# A new phased assembly of the Antarctic spiny plunderfish provides novel insights into the evolution of the notothenioid radiation

**DOI:** 10.64898/2026.04.21.719633

**Authors:** Jacopo Martelossi, Ksenia Krasheninnikova, Amy Denton, Jonathan M. D. Wood, Thomas Mathers, Richard Durbin, Nova Fong, David L. Bentley, Melody S. Clark, Iliana Bista

## Abstract

**Background:** Notothenioids are a well characterised species flock endemic to the Antarctic and an important model group for the study of genome adaptation to extreme cold. We used a new reference assembly and clade-wide comparative genomic analysis to investigate cryonotothenioid evolution and the appearance of novel functionalities linked to cold adaptation.

**Results:** A new phased assembly of a model notothenioid, *Harpagifer antarcticus,* demonstrated low levels of haplotypic variability across the genome. Nevertheless, numerous insertions from multiple LINE-L2 clades were found, suggesting ongoing transposition with potential contribution to speciation. Contrary to expectations the *afgp* locus was highly similar between haplotypes, except for large length allelic variants of *afgp* genes. Analysis suggests a model for the *afgp* locus expansion in *H. antarcticus* through segmental tandem duplications involving two pairs of *afgp* genes at time. Syntenic reconstruction of genomes from across the clade demonstrates conserved macrosyntenic relationships and group specific chromosomal fusions of notothenioids. Quantification of genome gain and transposition rates during cryonotothenioid diversification showed a first ancestral slow genome expansion concurrent with historic temperature drops. This was followed by lineage-specific massive peaks of genomic gain and transposition activity. Finally, we identified a set of genes that underwent ancestral diversifying selection and acquired novel conserved non-coding elements during the cryonotothenioid emergence. These were related to antioxidants and proteostasis, which may have facilitated the notothenioid Antarctic radiation.

**Conclusion:** Diversifying selection and genomic gain linked to transposon activity are primary contributors to lineage-specific evolutionary dynamics through the clade which facilitated adaptation to life in the cold.

## Background

The notothenioids are one of the most diverse and dominant fish groups in the Southern Ocean and a rare example of a well characterised marine flock species [1]. The clade is formed by approximately 140 species grouped in eight families, including three non-Antarctic (basal groups) and five mostly high specialised Antarctic families which are also known as cryonotothenioids [2]. They are characterised by genomic adaptations to extreme cold and they are considered to be an important model group for the study of cold adaptation [3].

The appearance of the antifreeze glycoproteins (AFGPs) is one of the most characteristic innovations enabling freeze avoidance in this group, and are present in members of the cryonotothenioid clade [4]. However, multiple other adaptations on the molecular and cellular level have also been considered to play important roles in the survival and establishment of notothenioids (e.g., cold stable enzyme production, lipid composition of cellular membranes, modified mechanisms of protein homeostasis) (summarised in [5]). The mechanisms through which such adaptations may have arisen, or the timing of their appearance along the ancestral cryonotothenioid clade remain poorly understood.

Assembly of notothenioid genomes has been confounded by the high number of repeat elements present in these genomes [6]. This is exemplified by the structure of the antifreeze gene locus, which is characterised by tandemly arranged genes [4,7], and high levels of repeat content making it very challenging to assemble and analyse accurately [6]. For the *afgp* gene locus in particular it has been suggested that high levels of structural variation on the haplotype level may be present. These estimates were based on BAC library sequencing reconstruction of the *afgp* locus of the Antarctic toothfish species *Dissostichus mawsoni* [8]. In that case it was shown that the two haplotypes of the *afgp* locus were characterised by extensive structural variation, with a distinct number of *afgp* gene copies, with a difference in overall size and mapping within each haplotype of the locus. Other reconstructions of the *afgp* locus based on whole genome sequencing data were only able to map the consensus structure of one assembled haplotype [6,9,10] though haplotig data have also been analysed [6]. To be able to test this hypothesis on other species, haplotype resolved (phased) genome assemblies are required.

Using Oxford Nanopore Technologies (ONT) and Hi-C data we assembled a new haplotype resolved chromosomal reference genome assembly for *Harpagifer antarcticus* (Antarctic spiny plunderfish). This is the first phased assembly for any notothenioid species. In addition, *H. antarcticus*, as a small, shallow-water and experimentally tractable member of the cryonotothenioids, is also an important model species for the study of cold adaptation [11–13].

Previous work on notothenioid genomes revealed the significant contributions of transposable elements (TEs) in the evolution of the clade. For the cryonotothenioid clade in particular it has been shown that a recent massive burst of transposons was linked to a significant genome expansion compared to the sub-Antarctic group [6,14,15], while the evolution of various other genomic features has also been linked to transposon activity such as expansion of the antifreeze locus, and the loss of haemoglobins in the Channichthyidae family [6]. Furthermore, it has been suggested that TE accumulation in some notothenioid groups, may have promoted karyotypic reorganisations likely leading to species diversification [16].

In general, TEs are known to be a major source of genomic variation across eukaryotes by disrupting gene structures, reshaping regulatory networks, and co-opting novel regulatory elements [17]. The extensive presence of TEs throughout the genome might amplify the occurrence of recombination events thus promoting deletions, inversions, and translocations [18], while the presence of environmental stressors may also promote TE activity [19]. Using whole genome comparisons can be a powerful method for understanding the mechanisms and patterns of transposon activity at a large scale, while it could also enable highly accurate quantification of genomic losses and gains occurring during diversification events [20–22].

In the present study we employed large scale genome wide analyses of chromosomal references across the notothenioid radiation along with novel genomic resources, to understand the 1) overall contribution of transposable elements in the cryonotothenioid diversification, through timed and quantified analysis of genomic gains and losses, 2) the evolutionary dynamics of the antifreeze gene locus across the cryonotothenioids, and models of expansion of *afgps* and haplotypic diversity in the *H. antarcticus* phased assembly, 3) impact of other genomic features that may have enabled genomic adaptation to extreme conditions.

## Results

### Genome assembly and gene annotation

The assembled genome for *H. antarcticus* using ONT long reads and Hi-C data resulted in a fully phased assembly with two high-quality haplotypes (**Fig. 1**, **Fig. S1**). The genome was assembled at a total length of 1,069 Mb (Hap1) and 1,222 Mb (Hap2), and contig N50 of 17 Mb (Hap1) and 12,8 Mb (Hap 2). Each haplotype was assigned to 24 chromosomes (**Fig. 1A, C**) with a scaffold N50 of 44.8 Mb (Hap1) and 41 Mb (Hap2) (**Table S1)**. Both haplotypes were highly complete, with 99% BUSCO completeness for both haplotypes, and QV scores of QV 40 (Hap 1) and QV 40.4 (Hap2) (**Table S1**). Chromosomes were named based on syntenic relationships with *Cottoperca gobio* genome assembly (fCotGob3.1) [23] (**Fig. 1C; Table S2**). Completeness of the assembly was confirmed by a clear enrichment of telomeric repeats on at least one end of all chromosomes on both haplotypes (**Fig. 1A, Fig. S2)**, confirming previous higher level localisation data using *in situ* hybridization data of the (TTAGGG)_n_ repeat motif [24]. Based on *k*-mer analysis, the heterozygosity rate of the *H. antarcticus* genome was estimated at 0.37%.

**Fig. 1:**
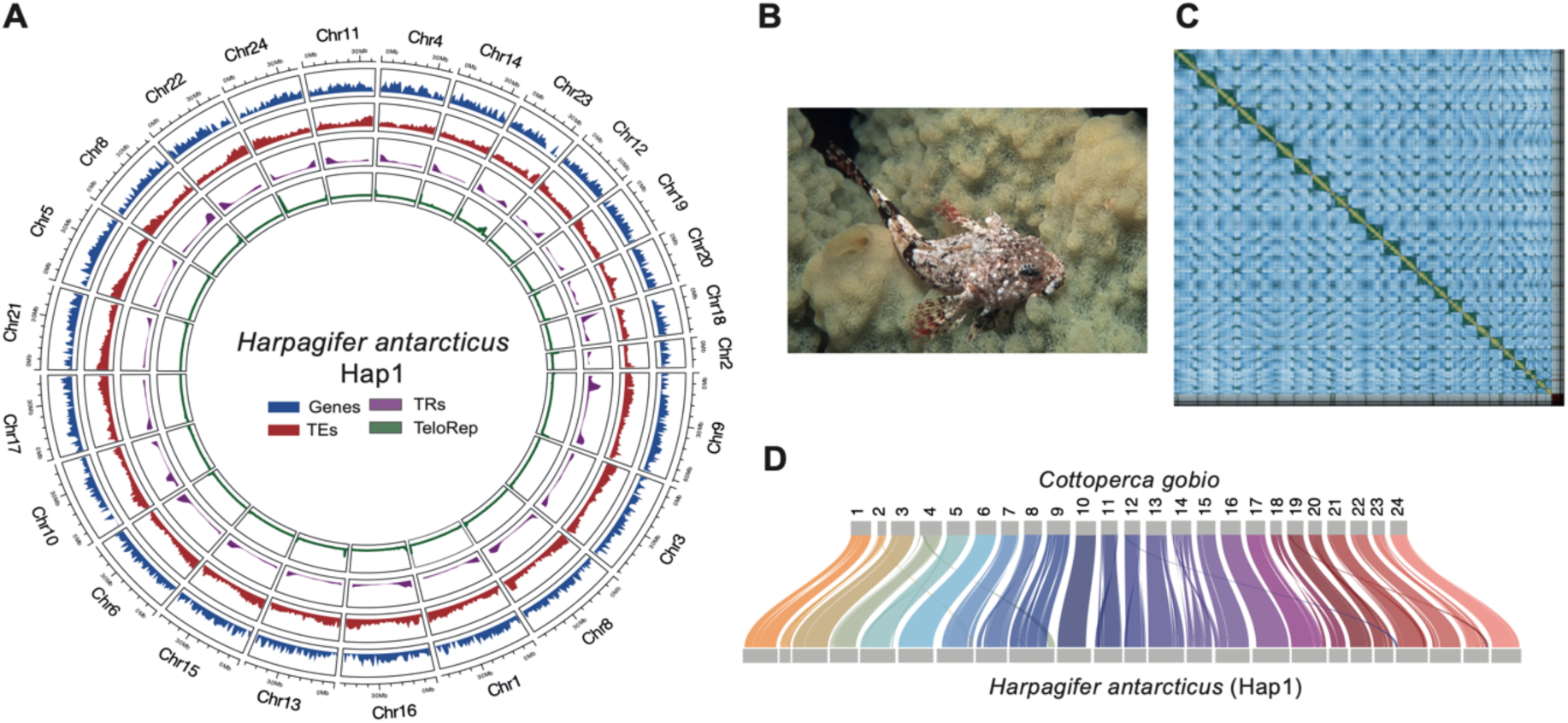
Genome assembly and genomic features. **A.** Circos plot showing the 24 chromosomes of the fHarAnt1.2 genome assembly (Hap1) and the distribution of protein coding genes (Genes), transposable elements (TEs), tandem repeats (TR), and canonical telomeric repeats (TeloRep). Chromosomes are ordered by length and named based on syntenic positions with *Cottoperca gobio* reference genome [23]. **B.** Photograph of a *H. antarcticus* specimen from Ryder Bay, Antarctic Peninsula (Photo credit: BAS Photo Library. Photographer: Simon Brockington). **C.** Hi-C-contact map of fHarAnt1.2 Hap1. D) Syntenic relationships between *H. antarcticus* Hap1 and *C. gobio g*enomes.

De-novo gene annotation (BREAKER) for Hap1 identified 23,057 protein-coding genes, with an average length of 14,904 bp, an average of 8.3 introns per gene, and mean intron and exon lengths of 1,555 bp and 174 bp, respectively (**Table S3**). By lifting Hap1 annotation onto Hap2 we identified 22,463 (**Table S3)** protein coding genes with 696 flagged as pseudogenes (i.e., containing premature stop codons). BUSCO completeness of the gene annotation was high for both haplotypes with 97.7% for Hap1 and 95.6% for Hap2 (**Table S3**).

### Genomic landscape of repetitive elements

Annotation of transposable elements on the fHarAnt1.2 genome revealed 41% coverage (Total repeats 43%) (**Table S4**) with a clear enrichment towards chromosome ends (**Fig. 1A**). Each of the three major TE classes (LINEs, LTRs, DNA), were almost equally represented in Hap1, covering approximately 13% of the genome respectively, whereas LTRs are more prevalent on Hap2 (17.84% of the genome). Classification at the RepeatMasker superfamily level was achieved for just 32% of LTR and 41% of DNA elements, in contrast to 97% of LINE elements. These predominantly consisted of LINE-L2 which accounted for 9.6% (Hap1) and 8.44% (Hap2) of the genome (**Table S4**). Tandem repeats and simple repeats accounted for about 4.5% and 2% of the genome on the haplotypes, respectively. Repeat landscape profiles describing activity of TEs through absolute time revealed that the TE content of the Antarctic spiny plunderfish genome mainly derived from a recent burst of activity of all three major TE groups that occurred after the diversification of cryonotothenioids, between 4 and 7 Million Years Ago (MYA) (**Fig. 2A**).

**Fig. 2:**
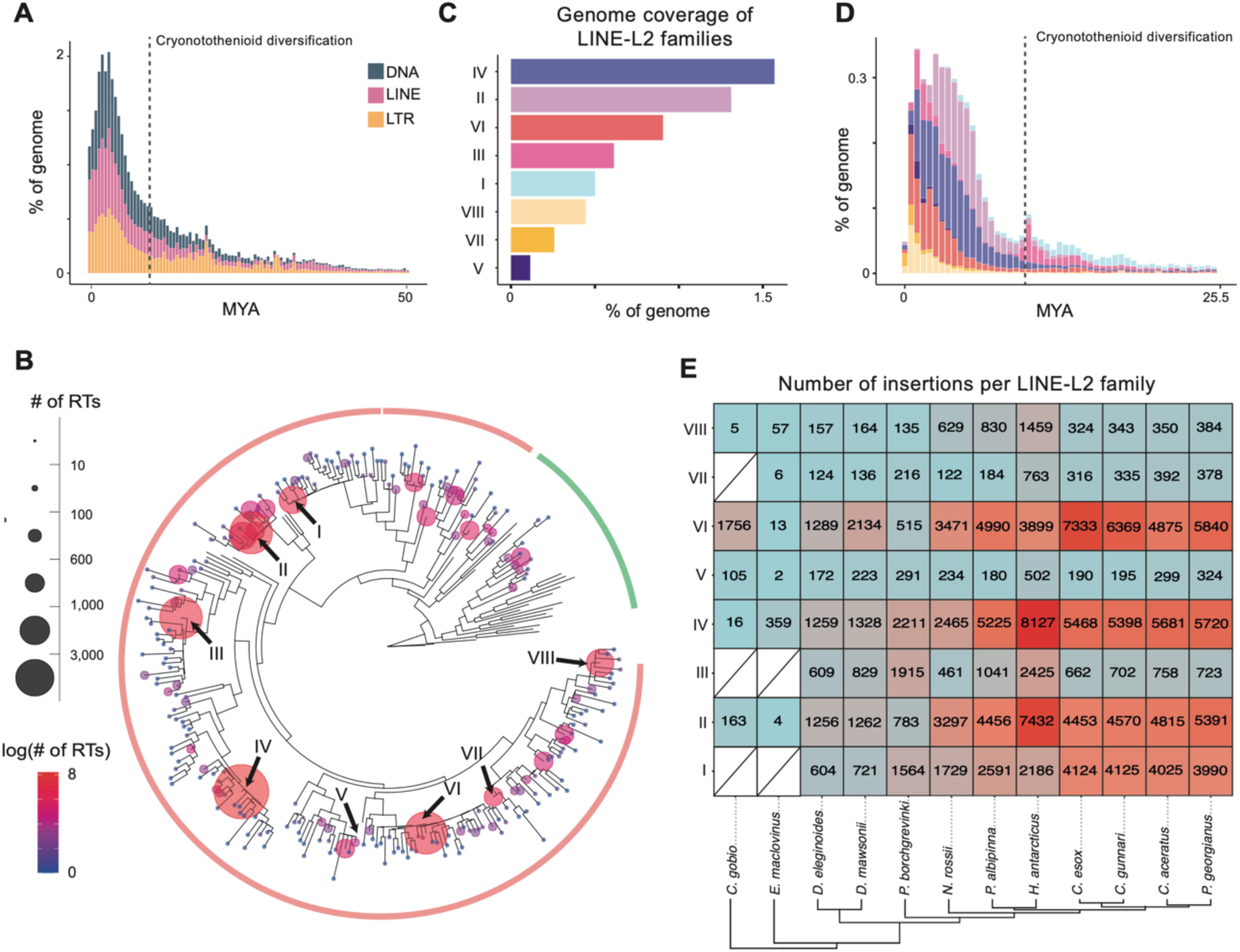
LINE-L2 elements in the *H. antarcticus* genome. **A**. Repeat landscape profile showing the activity of the three main TE groups through absolute time. Each bin represents one million years (MYA = million years ago). **B**. Maximum likelihood phylogenetic tree of clustered LINE-L2 reverse-transcriptase sequences (fHarAnt1.2 genome). Tips represent the longest element for each cluster. The size and colour of bubbles is proportional to the number of elements contained in each cluster. Arrows highlight the eight identified high-copy number families. Tips without bubbles represent reference sequences extracted from [25]. **C**. Genome coverage per LINE-L2 family, and **D**. activity through absolute time of the eight LINE-L2 families highlighted in panel B. Each bin represents 500 thousand years. **E**. Number of insertions across notothenioid genomes for each LINE-L2 family identified in *H. antarcticus*.

To further characterize the evolutionary history of the highly abundant LINE-L2 elements we first employed a phylogenetic approach to identify evolutionarily distinct LINE-L2 families (**Fig. 2B**). Among the 33,670 insertions harbouring a reverse transcriptase segment, 86% clustered with reference L2 elements, whereases others are part of other L2-related clades (Crack, Daphne L2A and L2B) [25]. Eight different high copy-number LINE-L2 families were identified, which were named HarAnt_LINE-L2_I – VIII. These families accounted for 60% of the total LINE-L2 content, and 57% of the genome (both haplotypes). The two most abundant of these were L2_IV (27.4% LINE-L2, 1.5% of the genome), and L2_II (22.9% LINE-L2,1.3% of the genome) (**Fig. 2C).** Dated repeat landscape plots suggest that all families were already active before the extant cryonotothenioid radiation, with a first peak concurrent with their emergence time ∼10 MYA and a second more recent activity peak between 4 MYA and 500 thousand years ago (**Fig. 2D**). The two most abundant appear to have decreased their transposition rate over the last 1 MYA, whereas the less abundant ones LINE-L2_VI, LINE-L2_VII, and LINE L2_VIII, have become predominant.

Subsequent analysis of LINE-L2 families across 12 notothenioid genomes (**Table S5**), identified all LINE-L2 families across all cryonotothenioid species. The three most abundant LINE-L2 families in the *H. antarcticu*s genome were also the most abundant ones in the other species (**Fig. 2E**). In the sub-Antarctic species *E. maclovinus* and *C. gobio*, some low-abundance families, 2 and 3 respectively, were missing and therefore likely have originated along the cryonotothenioid stem branch (**Fig. 2E**).

A total of 170 (18S), 534 (5S), 166 (5.8S), and 181 (28S) ribosomal RNA (rDNA) genes were annotated in the *H. antarcticus* genome. Among the 18S, 5.8S, and 28S genes, multiple copies were found to be dispersed throughout the genome, however, the majority, 97.1% of 18S, 97.6% of 5.8S, and 97.2% of 28S, were located in a single main 45S rDNA locus at the end of chromosome 18 (**Fig. S3 A, B**). The 5S genes occur in three distinct contexts: (i) within the 45S locus forming 18S–5.8S–28S–5S arrays (155 copies), (ii) as 20 interspersed single copies scattered across the genome, and (iii) in two large tandem arrays on chromosomes 13 and 21 (**Fig. S3 A, C**), which contain 151 and 208 paralogues, respectively. These data expand our knowledge of 45S and 5S rDNAs in *H. antarcticus*, which were previously thought to be co-localised to a single locus [26]. However, the dispersed nature of the 5S locus is consistent with reports for other notothenioids and fish in general [26].

### Haplotypic variation in the *H. antarcticus* genome

Alignment of the two haplotypes of the fHarAnt.1.2 assembly showed heterozygosity rate of 0.15%, consistent with the low heterozygosity estimated based on *k*-mers, and a generally low number of structural variants (**Table S6; Fig. S4**). More specifically 1.68% of the genome was present in a hemizygous state (i.e., composed of heterozygous insertions and deletions), and 8.9 Mb and 9.1 Mb was affected by heterozygous inversions on Hap1 and Hap2, respectively. Hemizygous deletions relative to Hap1 affect a total of 849 exons, and 136 genes are completely deleted on Hap2 and thus might be subjected to presence-absence variation at the population level. Importantly, 66.6% of all SVs were also detected in a heterozygous state when performing a read-based SV call, while only 1.7% were genotyped as homozygous for the alternate allele, indicating low levels of false positives due to assembly errors. The remaining 31.7% were not detected using ONT reads. Using generalized linear mixed models and chromosome identity as random effect, a significant positive association was observed between SV occurrence (considering inversions, duplications, and translocations) and repetitive content within 50 kb flanking regions of the breakpoints. The strongest predictor of SV occurrence was the density of tandem repeats, with an odd ration of 2.7. Additionally, we found a negative association of SV occurrence with gene content, distance from both chromosome ends, and assembly gaps as well as LINE-L2 content (**Fig. 3A, B**).

**Fig. 3:**
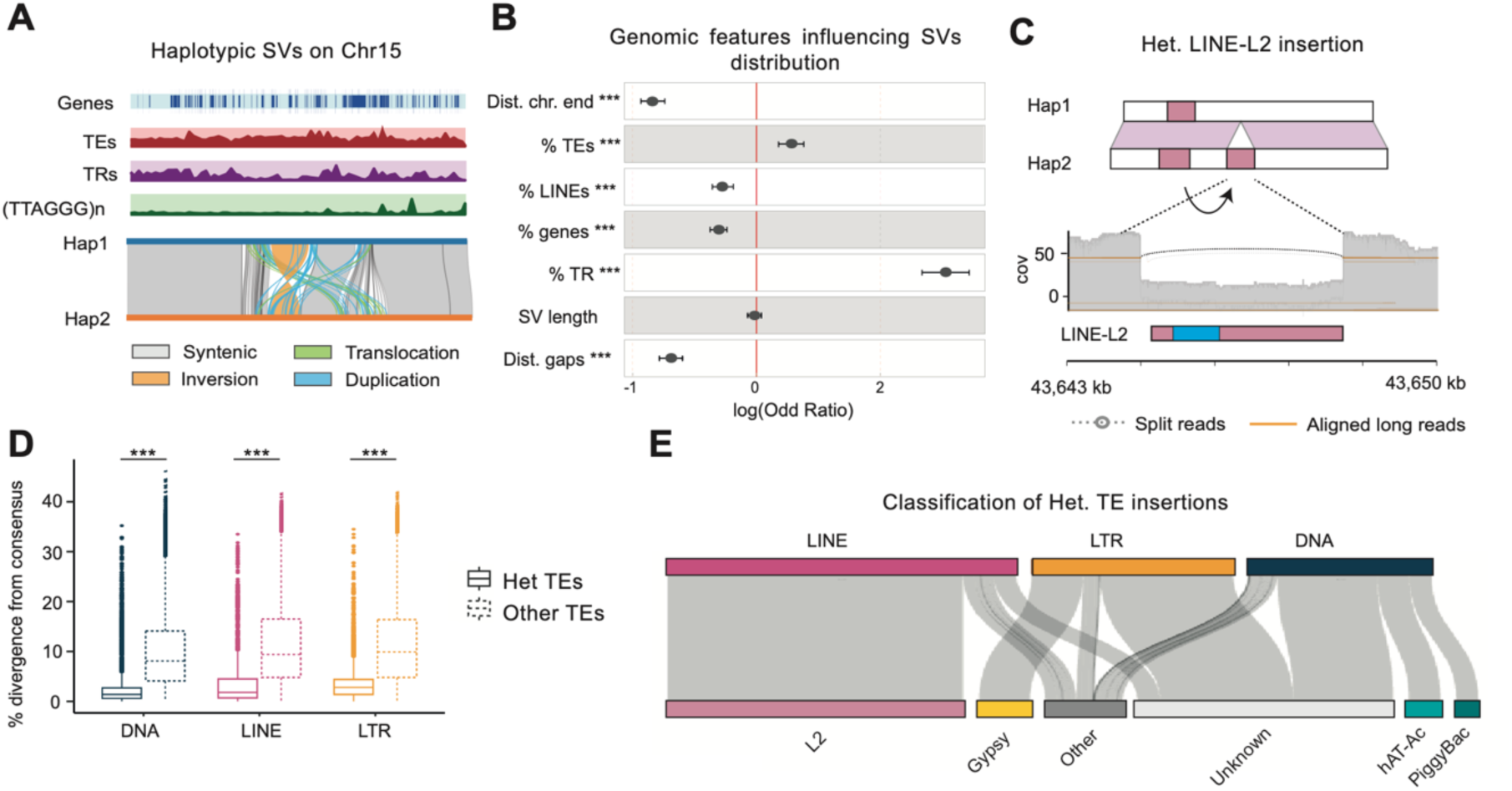
Haplotypic variation in the *H. antarcticus* genome. **A.** Detail of the right end of chromosome 15 harbouring multiple structural variants (SVs). **B.** Association between different genomic features and SV distribution, using a randomized control expressed as log-odds ratios. Points represent the estimated coefficients from the mixed-effects logistic regression. Horizontal lines indicate 95% confidence intervals (“Dist. Chr. end”: distance from closest chromosome end “Dist. Gaps” distance from closest genomic gap percentage (%) of TEs, LINEs, genes, and tandem repeats shown along 50 kb genomic regions flanking the SV breakpoints, SV: length of SVs). **C**. Schematic representation of a heterozygous LINE insertion between haplotypes. The blue rectangle marks the reverse transcriptase domain. **D**. Percentage of divergence of TEs from their source consensus sequences as a proxy of the time of the insertion of heterozygous, and all other TEs. **E**. Contribution of TE groups to heterozygous insertions (see detailed figure in **Fig. S5**) (*** = p-value <0.001).

The phased assembly allowed the inspection of the genome for the detection of potentially recently active TEs, as recent insertions are expected to be found mostly in a heterozygous state and thus correspond to insertion/deletion events between the two haplotypes (**Fig. 3C)**. Overall, 20% of the insertions/deletions have a reciprocal overlap of at least 75% with annotated TEs and might therefore correspond to heterozygous TE insertions. Accordingly, these TEs have a significantly lower divergence from their consensus sequences than other TEs, which is used as a proxy for insertion time (One-tail Wilcoxon rank sum test, p-value <0.001) (**Fig. 3D)**. A 39.1% of all heterozygous TE insertions correspond to LINE-L2 elements followed by unknown LTRs with 17.2%, and unknown DNA transposons with 12.4% (**Fig. 3E; Fig. S5**). Among all heterozygous TE insertions, most are located in intergenic regions (9,578), followed by 2,500 bp gene flanking regions (3,800) and introns (1,958). Only 23 insertions affect coding exons, leading to allelic pseudogenization of the impacted genes.

### Genome evolution during notothenioid diversification

Synteny analyses of 12 notothenioid chromosome-scale genomes (**Fig. 4A; Table S5**) revealed strong conservation of macrosyntenic relationships during their diversification, together with lineage-specific Robertsonian chromosomal fusions (**Fig. 4B**), in line with previous karyotype analyses [27]. Most assemblies, including *H. antarcticus*, show the predominant notothenioid karyotype of n=24, which largely comprises acrocentric chromosomes with the exception of *Notothenia rossii* (Nototheniinae) (n=12), and *Pagothenia borchgrevinki* (Trematominae) and *Pogonophryne albipinna* (Artedidraconidae) (n=23). In *N. rossi* 10 of the chromosomes appear to have originated from the Robertsonian fusion of 10 ancestral acrocentric pairs, one from the fusion of three chromosomes, and one remained unfused leading to a karyotype of *n*=12. In *P. borchgrevinki* and *P. albipinna*, only two different chromosomes have been fused, resulting in *n*=23 karyotype in both species. Across all notothenioid genomes, detailed molecular mapping detected only a few large inversions and intrachromosomal translocations were detected, mostly at chromosome ends (**Fig. 4B; Fig. S4**).

**Fig. 4:**
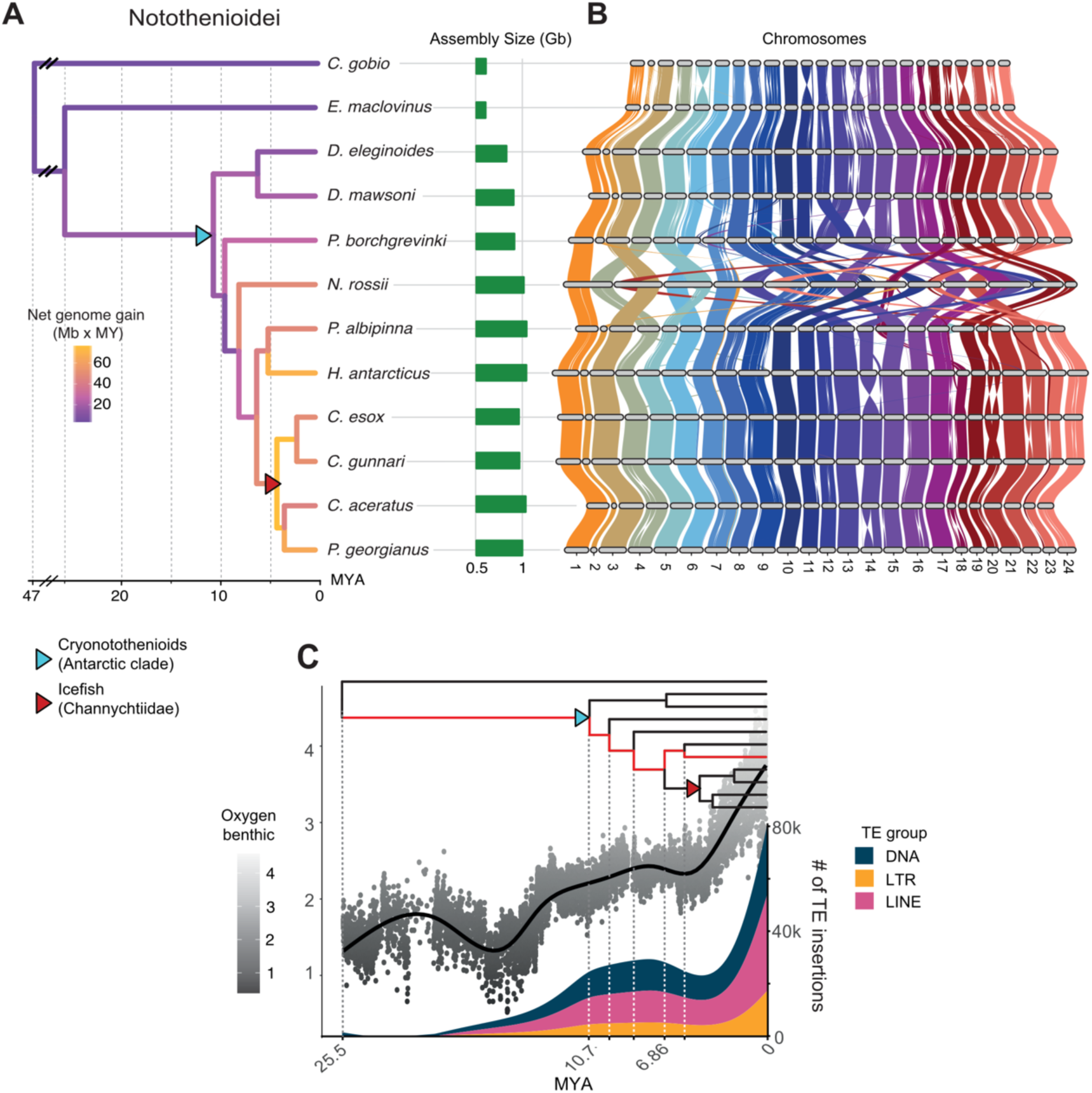
Genome evolutionary dynamics during notothenioid diversification. **A.** Dated phylogeny of notothenioid species included in all comparative genomic analyses. Branch colours show the rate of net genomic gain per million years as megabases per million years (Mb x MY). **B.** Syntenic relationships among notothenioid chromosomes. Chromosomes were named based on synteny with *C. gobio* assembly. **C.** Number of TE-related insertions along *H. antarcticus* ancestral and terminal branches (highlighted in red in the phylogenetic tree above), shown as a smoothed curve. Scatterplot and black curve show temperature variation over time based on deep-sea δ^18^O records (data from [28]).

We used a whole genome alignment of 12 chromosomal assemblies to understand patterns of DNA gain and loss during notothenioid diversification, and quantified the amount of bp involved in insertion and deletion events across all internal and terminal branches. Overall, genomic gain greatly outweighs genomic loss which accelerates towards recent times, reaching a maximum net rate of 76 Mb per MY along the ancestor of the Channichthyidae family (**Fig. 4A; Table S7**). Along the cryonotothenioid stem branch we observed a net genome expansion of 256 Mb. Although this represents the fourth largest absolute gain across all branches, surpassed only by the terminal expansions in *N. rossii*, *H. antarcticus*, and *P. borchgrevinki,* the long branching time resulted in a relatively modest rate of genome gain of 16.46 Mb per MY (**Fig. 4A; Table S7**).

Between 3,107 and 51,757 of the insertions inferred along *H. antarcticus* ancestors, map to an annotated TE element of its genome, requiring a reciprocal overlap of 75% between the two annotations and thus likely correspond to genuine TE insertions (**Table S8**). The number of TE insertions per MY greatly increased during early evolution of cryonotothenioids, between 10.7 - 6.86 MYA, slightly decreased along the lineage leading to the *Harpagiferidae* and *Artedidraconidae* sister species, and reached a main spike of activity along *H. antarcticus* terminal branch (10,456 insertions per MY; **Fig. 4C).** DNA transposons were the most active along all branches, but LINEs were predominant along the cryonotothenioid stem branch (41% DNA vs 42% LINEs) and, particularly (**Table S8**), *H. antarcticus* terminal branches (33% DNA vs 46% LINEs). Patterns of TE activity, especially for LINE elements, mirror variations in temperature data, based on δ^18^O variability [28] (**Fig. 4C**), with higher values related to lower temperatures.

### Conserved elements and selection analyses on protein-coding genes

We identified genes and non-coding regulatory elements that may have facilitated the successful colonization and diversification of cryonotothenioids in the cold waters of the Southern Ocean. First, we analysed intergenic conserved elements (CEs) which are arising along their stem branch and are conserved among all analysed species based on the notothenioid whole genome alignments, and second, we investigated signs of diversifying selection on protein coding genes along the same ancestral branch using a gene-wise branch-site model. We expect that genomic innovations underlying cold adaptation arose along the notothenioid ancestral lineage. Although some cryonotothenioids such as *D. eleginoides* and *C. esox*, inhabit more temperate regions, by colonizing Patagonia and several sub-Antarctic islands [29,30], these are nevertheless considered secondary adaptations [31]. Prediction of CEs yielded a total of 675,818 conserved genomic regions accounting for a total of 78,685,553 bp (**Fig. S6A**). Most CEs (78%) are shared by all notothenioids, whereases only 10,182 emerged in the cryonotothenioid ancestor and are shared by all members of the Antarctic clade (cryonotothenioid-specific CEs) (**Fig. S6**). Among the cryonotothenioid-specific CEs, most correspond to intergenic (42.9%) and exonic (29.3%) genomic regions. Intergenic, non-coding CEs (CNEs) might play a regulatory role in gene expression. We found that the 1,261 genes associated with at least one cryonotothenioid-specific CNE were enriched for a total of 236 Gene Ontology (GO) terms, most of which are related to signalling, cell differentiation and developmental processes (**Table S9**). However, we also identified 120 genes involved in biological functions known to be important for cold adaptation, in processes such as antioxidant activity, lipid metabolism, protein folding, and cytoskeleton organization (**Table S10**). Some genes of particular interest including *prdx5* associated with antioxidant activity (**Fig. S6B**), *ube2e1* and *ube2h*, involved in protein ubiquitination; *ahsa1* which has chaperone-binding activity, and *tubb4a*, with Antarctic fish tubulins previously having been identified as cold adapted at the amino acid level [32].

For selection analyses we used an extended dataset which includes an additional seven temperate or tropical fishes tagging on the cryonotothenioid stem branch (**Fig. S7**). To avoid conflating the signal with lineage-specific evolutionary dynamics, we did not test for diversifying selection in any cryonotothenioid child branch. Furthermore, to establish a stronger link between the identified genes and cold adaptation, we also excluded genes that were found to be under diversifying selection in any other non-cryonotothenioid temperate or tropical fish, based on a control analysis (**Fig. S7**). While we acknowledge that this latest approach might increase the false negative rates, excluding genes that might be related to both cold-adaptation and other phenotypic traits due to for example substitution on different codons [33] we prefer to automatically exclude genes that might be subjected to pervasive diversifying selection along multiple branches of the phylogeny. We identified a total of 334 genes under selection on the cryonotothenioid ancestor but not in temperate-adapted species, of which 256 were functionally annotated (**Table S11**). A total of 54 GO terms were enriched among these genes (**Table S12**). Of these, several are potentially linked to cold adaptation, particularly to oxidative stress (GO:0070301, GO:0010039), lipid metabolism (GO:0006084, GO:1903727, GO:0006690) and proteostasis (GO:0051260, GO:0051603, GO:0006509, GO:1901984). Example gene include the sirtuin 1 (*sirt1*), peroxiredoxin 3 (*prdx3*), mitochondrial superoxide dismutase 2 (*sod2),* and dihydrolipoamide dehydrogenase (*dld*) genes.

### Characterization and evolution of the antifreeze gene locus

The antifreeze gene locus was located on chromosome 16, spanning approximately 699 kb on each haplotype, in a region flanked by the *hsl* and *tomm40* genes. Using a combination of automated and manual approaches, we identified 17 and 16 *afgp* gene copies on Hap1 and Hap2, respectively, as well as two chimeric *afgp/tlp* genes on each haplotype (**Fig. 5A**). The gene annotation completely matched the *afgp*-specific k-mer signature identified by Klumpy suggesting a complete representation of the locus (**Fig. S8**). Repeat coverage of the *afgp* locus was 58.7% (Hap1) and 59.1% (Hap2), approximately 1.43 times higher than the genome-wide estimation (**Fig. 5A**).

**Figure 5:**
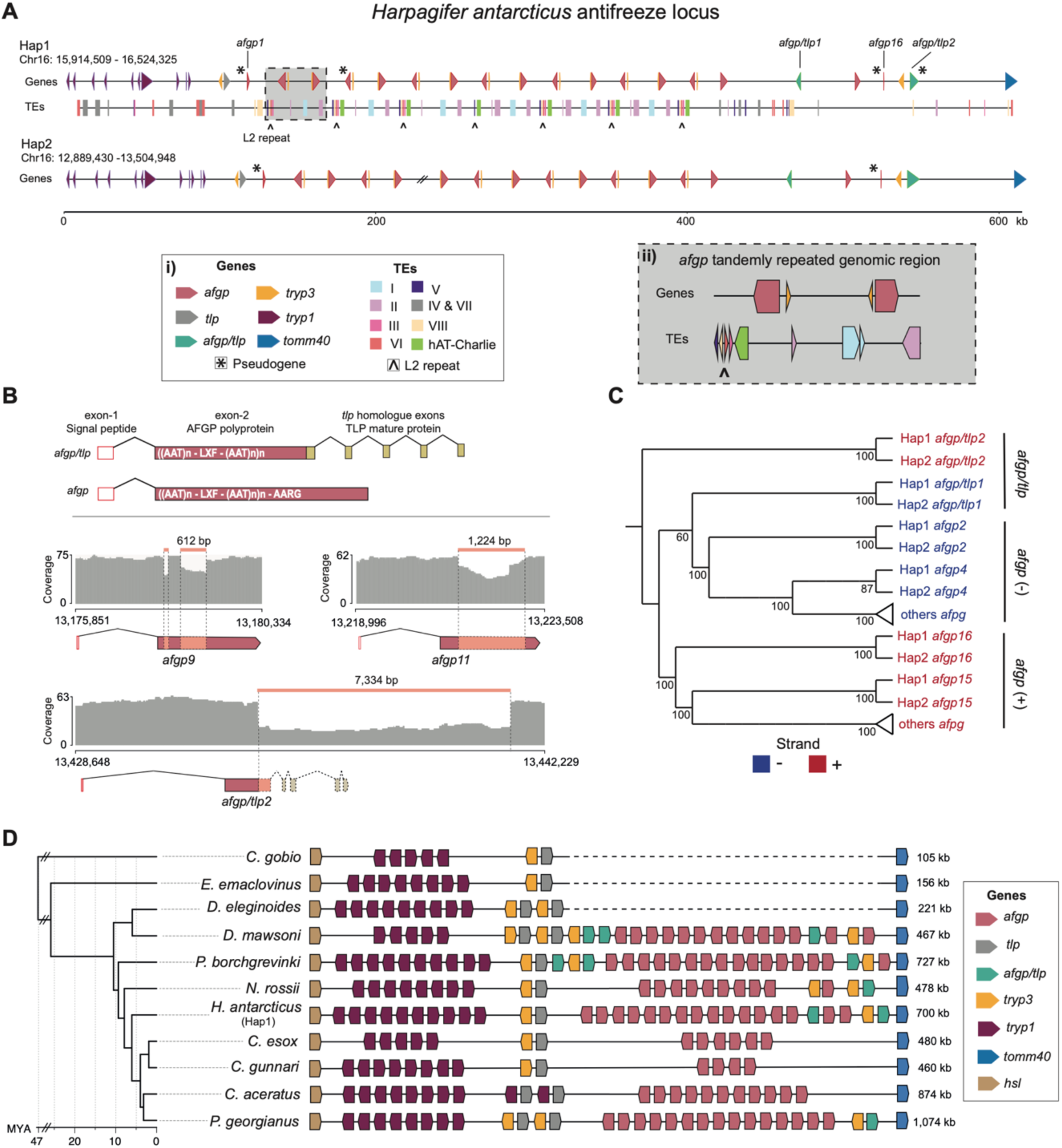
Structure and evolution of the antifreeze locus. **A.** Structure of the antifreeze gene locus on both *H. antarcticus* haplotypes. Triangles represent different genes, and rectangles represent transposable elements. *Afgp* genes are named from 1 to 16 based on their position on the locus. Asterisks highlight pseudogenes and double-slash lines indicate assembly gaps. The grey panel (ii) on the right corner shows a zoomed in version of the tandemly repeated genomic unit marked by the characteristic TE motif. **B.** Top: simplified representation of *afgp* and chimeric *afgp/tlp* genes (exon and intron lengths are not proportional to their actual size). Bottom: representative *afgp* and *afgp/tlp* chimeric genes subjected to insertion/deletion events between the two haplotypes. Coverage refers to ONT reads mapped back to assembly. **C.** Phylogenetic tree of *afgp* and chimeric *afgp/tlp* gene copies. Different colours correspond to strand (+/-). **D.** Structure of the *afgp* locus across notothenioids. All genes, including pseudogenised copies are reported. Locus for *P. borchgrevinki*, *C. esox*, *C. gunnari*, *C. aceratus*, *P. georgianus* and *D. mawsoni* is based on published work [6,8,9,34]. *P. albipinna* could not be included in this analysis due to the locus being highly fragmented in the corresponding assembly (**Fig. S8**).

The typical structure of a functional *afgp* gene encodes for an exon-1 signal peptide and an exon-2 AFGP polyprotein with XLF spacer and AARG terminal motif [4] (**Fig. 5B)**. On Hap1, 14 of the *afgp* genes identified contained the correct structure and were thus considered to be functional. All complete *afgp* copies except one were automatically predicted by GeMoMa, highlighting its strong ability to detect intact *afgp* genes (**Fig. S8**). Three more identified copies were putative pseudogenes (**Table S13**), two of which miss the exon-l signal peptide (*afgp1* and *afgp17*) (**Fig. 5A**). The third copy (*afgp4*) encodes a partial AFGP polyprotein with an unrelated amino acid tail sequence that lacks the canonical AARG terminal motif. The two chimeric *afgp/tlp* genes contain a signal peptide, but while *afgp/tlp1* encodes the XLF spacer and a short TLP segment, *afgp/tlp2* encodes an unrelated amino acid tail sequence and lacks the XLF spacer, making it a putative pseudogene (**Fig. S9**).

The structure of the *afgp* locus of fHarAnt1.2 is almost identical in both haplotypes, without evidence of major rearrangements. One *afgp* gene copy is missing in Hap2 (*afgp6*) compared to Hap1, but this may be likely due to a mis-assembly resulting in a gap (**Fig. 5A**). Only insertions and deletions were identified between the two haplotypes, with 48 short variants (<50bp), involving a total of 428 bp and other 16 with a length from 51 bp to 7,477 bp affecting a total of 32,109 bp. The previously mentioned pseudogenization of *afgp4* on Hap1 resulted from a single-base deletion that shifts the reading frame relative to Hap2, which retains a functional copy. Here, 22 insertions/deletions occur on exon2 of *afgp* or *afgp/tlp* genes affecting a total of 3,27 kb. These variants do not disrupt the reading frame but generate length allelic variants of the AFGP polyprotein. Specifically, we identified five large indels with a length between 51 bp and 1,224 bp affecting a large portion of exon-2 of *afpg7, afgp9* and *afgp11* genes, respectively (**Fig. 5B)**. Finally, the chimeric *afgp/tlp2* gene on Hap1 is affected by the largest identified SV within the locus: a large deletion of 7,334 bp that involves all *tlp*-derived exons encoding the TLP mature protein (**Fig. 5B**).

The locus is characterized by a clear and repetitive organization with 14 *afgp* gene copies found in an area spanning 287,044 bp. The genes and TEs in this region are arranged in a characteristic motif consisting of two pairs of *afgp* genes with alternating orientation, and a specific set of TE insertions (**Fig. 5Aii**). The motif is consistently repeated along the region suggesting that it mainly consists of segmental duplications (**Fig. 5A**). Mapping of ONT, and 10X reads (minimum mapping quality >0, and excluding secondary alignments) from [6] which come from the same individual, revealed sharply reduced coverage within the previously mentioned central region of the locus (**Fig. S10 A, B, C**). On average, coverage decreased by 16.8% for ONT reads and by 55.5% for 10X reads. These values reach the 85.4% and 32.8% when only reads with a minimum mapping quality of 20 are considered. Under this threshold 32.9% of the locus show no coverage of 10X reads.

To better understand the evolutionary dynamics and duplication history of *afgp* and *afgp/tlp* genes we inferred a maximum likelihood tree based on the conserved exon-1 and the starting region of intron-1 which is shared between *afgp* and chimeric genes (**Fig. S11; Fig. S12**). Due to the difficulties in resolving relationships among highly similar *afgp* gene copies, based on few alignment positions (391, of which 56 are parsimony informative) (**Fig. S12A**), we additionally performed a ML tree inference excluding chimeric genes (**Fig. S12B**). This allowed us to include the full intron-1 and analyse a total of 1,964 positions with 3.2 times more parsimony-informative sites. Because both trees recovered the same two main *afgp* clades, we combined the two topologies by placing the chimeric genes onto the *afgp*-only tree, preserving their phylogenetic positions (**Fig. 5C**). The chimeric *afgp/tlp2* genes from both haplotypes form a clade separated by a long branch from all other genes (**Fig. S12A**). Considering that *afgps* likely originated from a chimeric gene (Chen et al., 1997), *afgp/tlp2* probably represents the most ancestral gene and was therefore used to root the tree. All other analysed genes form two clades that perfectly separate genes on the plus strand from those on the minus strand, including the chimeric *afgp/tlp1*, which is also the first to diverge within its clade.

To have a broader overview of the locus across cryonotothenioids we also annotated the locus in additional species. We used an automated re-annotation process of the *afgp* locus by employing GeMoMa and Klumpy, followed by visual inspection of candidate genes. To benchmark this approach we used the *afgp* locus of species *P. georgianus* which had previously been manually annotated, which we then re-annotated through the automated process. The automated approach identified the same number of chimeric *afgp/tlp* (1) and *afgp* (15) genes in *P. georgianus*, organized in the same structure as described by [6]. We then auto-annotated two additional species: *P. albipinna* and *N. rossii* (**Fig. 5D; Fig. S8**). In *P. albipinna* the locus was distributed across different unlocalized scaffolds despite the use of long-read sequencing technologies and Hi-C data (**Fig. S8**). In *N. rossii* we detected two chimeric genes and 10 *afgp* genes. We performed comparison of the structure of the locus along a number of species including our new and published annotations for other included species (**Fig. 5D)**. Overall we detected some notable similarities between species *C. esox, C. gunnari, H. antarcticus*, and *N. rossii.* Specifically, in all these species the majority of *afgp* genes are arranged in arrays with alternating orientation. However, each pair of *afgp* genes lacks the previously described TE motif that was identified in the *H. antarcticus afgp* locus.

## Discussion

### Haplotype-resolved assembly of the Antarctic spiny plunderfish genome reveals low haplotypic structural variation

Here, we present the first haplotype-resolved genome assembly of a notothenioid species and one of the most contiguous notothenioid genome assemblies to date. The assembly was generated with ONT reads which supported assembly more successfully compared to sequencing attempts with HiFi data. The two haplotypes exhibit low levels of divergence and a relatively small number of structural variations, which is consistent with the low *k-*mer based and SNP based heterozygosity (0.37% and 0.15%, respectively) of the *H. antarcticus* genome (**Table S6**; **Fig. S4**). Few exons and genes were found to be absent from one of the two haplotypes, in contrast to findings in other metazoans, where highly divergent and hemizygous genomic regions can represent more than 50% of a diploid genome [35,36].

Complex SVs, such as translocations, duplications, and inversions were found to colocalize towards the repeat-rich and gene-poor chromosome ends (**Fig. 2A, Fig. S4**). This was even more so in terms of tandem repeat content, consistent with a model of elevated mutation rates in sub-telomeric genomic regions [37]. Accumulation of repeats towards chromosome ends was also observed in another cryonotothenioid species, *T. borchgrevinki* [34], so we suggest that this might be a plesiomorphic feature of the clade. Furthermore, recent TE insertions were found to significantly contribute to the emergence of insertions and deletions between haplotypes. Even though heterozygous TE insertions are usually found in intergenic genomic regions likely due to strong negative selection against TE insertion within genes, as observed in zebrafish [38], here a high number of TE insertions were detected in gene-flanking regions. In few cases, insertions within the exons of protein-coding genes were detected, which have led to the truncation and putative pseudogenization of the affected genes on one of the haplotypes. Overall, our results suggest that despite the low levels of overall haplotypic structural variability, recently active TEs likely represent important contributors in driving population-level differentiation in *H. antarcticus*, promoting gene-loss and reshaping *cis-* and *trans-* regulatory elements.

Furthermore, haplotype resolved assemblies would also have the potential to shed new light on overall genome evolutionary dynamics, facilitating assembly-based variant calling [39], pangenome reconstruction [40], allele-specific expression analyses [41], and TE identification [42].

### Recent accumulation of ancient LINE-L2 elements in *H. antarcticus*

Different TE clades can have varying evolutionary trajectories across species, reflecting host–element co-evolution, lineage-specific regulatory constraints, and variation in genomic defence mechanisms [43,44]. LINE-L2 transposons were previously shown to be particularly abundant in *H. antarcticus* compared to other notothenioids, covering about 8% of the genome [6]. Analysis of the new genome assembly similarly showed high LINE-L2 levels deriving mainly from the activity of eight different LINE-L2 families. Both the high number of heterozygous LINE-L2 insertions and the repeat-divergence landscape analyses suggest a recent, lineage-specific accumulation of these elements (**Fig. 2**). Eight evolutionary distinct families of LINE-L2 elements were identified, with concurrent activity within the last four million years (MY) (**Fig. 2D**). Moreover, most of these LINE-L2 families, including the most abundant ones, are also shared with sister species of the cryonotothenioid clade, suggesting their ancient emergence. Despite the recent activity of LINE-L2 elements, we observe a negative effect of their genomic content around breakpoint regions and occurrence of inversions, duplications and translocations between haplotypes suggesting that their accumulation has not resulted in the emergence of other more complex structural variants.

In another LINE transposon family, the LINE-L1 retrotransposons, it has been shown that either a single lineage was usually active at high replication rate at any given time in mammals, or multiple, ancient LINE-L1 lineages were concurrently active in fishes, reptiles, and bivalves [44,45]. The evolutionary path of LINE-L2s in notothenioids appear therefore to follow the generally observed evolution of LINE-L1 of fishes.

### Rampant genomic gain and low levels of genomic deletion during cryonotothenioid diversification

TE-mediated genome expansion has been shown to predate the diversification of extant cryonotothenioids and that enhanced TE activity continued in more derived clades such as icefishes [6,34]. To further test this hypothesis, we have quantified genomic gains through TEs along the notothenioid ancestral branches (**Fig. 4A**). Mapped TE insertions along *H. antarcticus* ancestral branches as well as a dated repeat landscape plot, indicate that most TE activity occurred recently, concurrently with the recent cryonotothenioid diversification. A single massive burst of TE activity was also detected in repeat profiles of other cryonotothenioids [6,15,34]. Low levels of ancient TE activity followed by recent peaks could reflect either genuinely increased transposition rates or extensive genomic deletions that removed older insertions, thereby erasing evidence of ancient bursts. However, net genome-gain estimates reveal low deletion rates during notothenioid evolution, supporting the former scenario: a slow accumulation of TEs before the emergence of extant cryonotothenioids, followed by a more recent increased rate of TE accumulation. Our reconstruction also revealed an ancient burst of genome expansion in the ancestor of icefishes explaining their exceptionally large genomes, which may be explained by sea temperature history in the region [6].

Paleoclimatic based temperature calculations (**Fig. 4C**) indicate an initial temperature drop at approximately 16 MYA which predates the emergence time of the extant cryonotothenioid clade, and a second, stronger reduction around 3 MYA which could have contributed to their recent diversification [6], similarly to other marine clades [46]. Here we show that the timing of these temperature variations appear to coincide with our reconstructions of TE levels of activity (**Fig. 4C**). LINE elements were found to be particularly expanded not only in *H. antarcticus* but also in other cryonotothenioids (**Fig. 2E**). The presence of environment induced stressors, such as fluctuations of temperature, have been linked to increased TE activity due to likely disrupting mechanisms of epigenetic regulation [47]. In the cryonotothenioid species *D. mawsoni*, it has been shown that the levels of LINE element transposition are increased through cold induced cell transfection [48]. We provide evidence that compared to other TEs, LINEs have been particularly active along the cryonotothenioid ancestor and starting from 4 MYA, concurrently with the two main temperature drops documented in the region (**Fig. 4C**). Overall, our comparative genomics analysis suggests that the increased genome size of cryonotothenioids may have occurred through a two-step process: one ancient, slow TE activity in the absence of major genomic deletions that predate the emergence of extant species and one more recent, lineage-specific massive bursts of transposition due to loss of epigenetic control.

### Macrosyntenic relationships and chromosomal fusions in notothenioid genomes

Our synteny analyses of 12 chromosome-scale notothenioid genomes demonstrated conserved marcosyntenic relationships between notothenioid chromosomes in accordance with the presence of a stable karyotype with low structural divergence during notothenioid radiation [49]. In the Nototheniidae, the most common chromosome number is 2n=48, mainly comprising acrocentric chromosomes. Indeed *H. antarcticus* has a karyotype of 2n=48 comprising 42 acrocentric, 2 metacentric and 4 submetacentric chromosomes [49]. Lineage-specific Robertsonian chromosomal fusions have occurred in some notothenioids [50], with the most notable case being *N. rossii* where only 12 chromosomes (2n=24) are observed (**Fig. 4B**). The few other identified smaller scale rearrangements tended to colocalize towards repeat-rich chromosome ends, similar to haplotypic structural variants identified in the *H. antarcticus* genome. These genomic regions are therefore thought to be subjected to higher rates of structural variation compared to the rest of the chromosomes, with repetitive elements that may drive this variability. Furthermore, the existence of LTR retrotransposon–insertion hotspots near chromosome ends and at chromosome fusion points, suggests that such repetitive sequences may have facilitated Robertsonian fusions, as has occurred in other more distant species [16,51]. In cryonotothenioids, reduction in chromosome number from the ancestral karyotype of n=24, caused by multiple independent chromosomal fusions, appears to be common within specific subfamilies, with such cases observed in the genera *Trematomus*, *Notothenia*, and *Pogonophryne* [24].

### Tandem segmental duplications and haplotype length variants shape the structure of the antifreeze locus

The antifreeze locus is a very challenging genomic region to assemble both because of its repetitive structure due to the presence of tandemly duplicated gene families such as the *afgps*, and trypsinogens, as well as due to its high TE content [6]. Using ONT reads we were able to assemble and annotate the complete *afgp* locus for *H. antarcticus* for both distinct haplotypes. The locus comprises 17 *afgp* copies (Hap1) and a higher repeat content compared to the genome average (59% vs 43%). One copy of the chimeric *afgp/tlp* gene was identified on each haplotype (**Fig. 5A**), both of which are characterized by canonical structure in accordance to previously described genes (Chen et al., 1997; Cheng and Chen, 1999; Nicodemus-Johnson et al., 2011) and are considered to be functional copies. These chimeric genes were previously considered to be putative evolutionary intermediates of the *afgp* genes [7]. Annotation of the locus on members of the icefish family showed either lack (*C. gunnari* and *C. aceratus*) or pseudogenisation (*P. georgianus*) of chimeric gene copies, suggesting that chimeric *afgp/tlp* genes might have been lost in low latitudes of the Southern Ocean by relaxed selection due to redundancy function with *afgp* genes [6,9]. However, chimeric genes have been retained in *H. antarcticus* which has a partially overlapping distribution range with these species, leading us to question the drivers of retaining these genes for this species.

BAC library-based analyses of the *afgp* locus of *D. mawsoni* suggested the presence of massive variation between haplotypes in both the overall size of the locus, and in the number of *afgp* gene copies [8]. Tandem arrays are generally expected to be frequently subjected to copy number variation due to recombination between homologue chromosomal regions which could also promote high levels of haplotypic variation [52]. Through this haplotype resolved assembly for *H. antarcticus* we were able to directly investigate the potential presence of haplotypic divergence in the locus. Nevertheless, we found that in *H. antarcticus*, both haplotypes exhibit almost identical structure and gene copy number. The main structural variations identified where pseudogenizations of one *afgp* gene due to a single base-pair deletion and of the only putatively functional *afgp/tlp* gene due to large deletion of all *tlp* derived exons on Hap1.

Despite low levels of variation at gene copy and functionality levels, we also identified allelic variation of the *afgp* gene copies, with regards to the actual length of the AFGP polyprotein coding exons. Potential variation on the length of the AFGP polyprotein could have important biological implications. The notothenioid AFGP proteins function through an “adsorption–inhibition” mechanism [53], adsorbing to the surface of ice and preventing the growth of the crystals as a function of glycoprotein concentration, size, and shape [54,55]. It has been observed that smaller protein molecules are less effective than large ones to interfere with ice crystal growth [54]. We could hypothesise that allelic variation linked to the size of AFGP polyproteins in *H. antarcticus* populations could represent an important source of genomic variation that allows local adaptation to more extreme conditions.

Length allelic variants have also been observed in other highly repetitive gene families, such as silk genes in caddisflies, butterflies, and spiders [56], where sequence length was linked to the properties of the silk fibres [57]. Such types of allelic variants may represent a convergence source of variation in proteins composed of repeated motifs across all metazoans. In these cases we could consider that because the gene itself provides a substrate for recombination, and the repetitive structure can weaken purifying selection against insertions and deletions, these could create a standing pool of genetic variation on which positive selection can eventually act.

In the case of the *afgp* locus, the characteristic TE plus *afgp* gene motif that was detected in *H. antarcticus* likely supports a specific mode of expansion of the locus. The *afgp* copies are arranged in opposite orientation separated into two strand-specific clusters (**Fig. 5A, C**), suggesting that the expansion of the genes occurred through tandem segmental duplications, which duplicated as two pairs of genes at a time. Furthermore, the high sequence homology identity across duplicated regions and the short branch lengths separating gene copies in phylogenetic analyses (**Fig. 5, Fig. S12**) suggest that these duplications are either evolutionarily recent or have been strongly homogenized by concerted evolution.

After their emergence in the cryonotothenioid ancestor, reduction and loss of *afpg* genes has occurred multiple independent times during their radiation [58]. However, the exact time at which *afgp* genes begun duplicating and expanding remains an open question. Comparison of the *afgp* locus structure across multiple notothenioid genomes (**Fig. 5D**) showed a similar genomic configuration of some genomic features in *P. borchgrevinki*, *H. antarcticus*, *C. esox, C. gunnari,* and *N. rossii*. Nevertheless, the distinct TE motif appears to be specific to the *H. antarcticus* genome (**Fig. 5Aii**). We hypothesize that the tandemly duplicated genomic region marked by the characteristic TE motif of *H. antarcticus* is a plesiomorphic feature of the other species, with different rates of duplication and loss driving variation of *afgp* gene copy number. We therefore suggest that from a hypothetical cryonotothenioid ancestral locus with few *afgp* genes, independent tandem duplications increased gene-copy number in different lineages.

### Diversifying selection and origin of novel conserved elements in cold-adaptive pathways during cryonotothenioid emergence

To detect additional genomic features underlying cold adaptation in notothenioids we searched for both protein-coding genes that experienced diversifying selection and CNEs that first emerged along their ancestral branch. Among the genes showing evidence of diversifying selection in the cryonotothenioid ancestor, many were found to be associated with biological processes known to be important for survival in extreme cold conditions (**Table S12)**. We found that most CNEs were shared across all notothenioid species, suggesting that a large portion of the genome expansion characterizing the cryonotothenioid ancestral lineage may have evolved neutrally. Additionally, most cryonotothenioid-specific CNEs were found to be associated with genes involved in signalling, cell differentiation, and developmental processes (**Table S9**). CNEs are known to have non-random distribution within the genome, because they have the tendency to cluster near genes with regulatory functions in multicellular development and differentiation (Polychronopoulos et al., 2017). A subset of the genes which we have found to contain newly acquired CNEs are related to two biological processes which are overrepresented among genes under diversifying selection. These biological processes are antioxidant activity and proteostasis and they have been linked to evolutionary cold adaptation [59,60]. We suggest that because these genomic innovations arose in the cryonotothenioid ancestor, they are amongst the key components of these pathways that enabled this clade to survive and become established in the Southern Ocean.

### Antioxidant activity and proteostasis

Life in freezing sea water poses particular biological problems for the organisms living there. The amount of oxygen increases considerably with reduced sea water temperatures, producing naturally high levels of cellular oxidative stress for Antarctic marine species [59,61]. In addition, cold denaturation of proteins impacts proteostasis [60]. Therefore, in order to survive in this harsh environment, organisms need to modify their biochemical processes.

Our analysis revealed four key genes related to oxidative stress that underwent diversifying selection in the cryonotothenioid ancestral lineage, including the sirtuin 1 (*sirt1*), peroxiredoxin 3 (*prdx3*), mitochondrial superoxide dismutase 2 (*sod2),* and dihydrolipoamide dehydrogenase (*dld*) genes. Moreover, we found that the peroxiredoxin 5 (*prdx5)* gene acquired novel CNEs concurrently with extant cryonotothenioid emergence. Gene *sirt*1 is a class III histone deacetylase that is recruited in the promoter region of multiple antioxidant genes, including *sod2, prdx3*, and *prdx5,* promoting their expression and their protein concentration [62]. These three genes encode mitochondrial enzymes that act in concert to control reactive oxygen species (ROS) levels. *Sod2* converts superoxide radicals into hydrogen peroxide, which is then detoxified to water by *prdx3* and *prdx5* [63]. The interplay between protein modifications and the emergence of novel, potentially regulatory CNEs within this pathway may have strengthened the antioxidant capacity of the cryonotothenioid ancestor, enabling it to cope with elevated oxidative stress in the mitochondria.

Signs of diversifying selection have been found [64] also in other cold-adapted notothenioids acting on genes involved in antioxidant activity, such as in *N. coriiceps* [65] and *D. mawsoni* [15], and across species. Moreover, genes such *sod2* and *prdx5* which are involved in ROS scavenging, have been found to be overexpressed in cold-adapted notothenioids compared to their temperate-adapted relatives [66]. Superoxide dismutase appears to be particularly important in combatting oxidative stress in Antarctic species. Gene duplications used to enhance the cellular activity of this enzyme have been identified in Antarctic invertebrates [67]. Furthermore, evolution to function efficiently in the cold have resulted in at least some members of this gene family being particularly thermally sensitive in some Antarctic marine invertebrates [68]. This would suggest that specific protein modifications might help to mitigate cellular oxidative stress in Antarctic species.

In terms of proteostasis, there is evidence that Antarctic species, including fish, face challenges in producing, folding and maintaining proteins at such low temperatures [60]. Whole animal RNA to protein ratios, ubiquitylation rates, and levels of chaperone proteins are much higher in Antarctic compared to temperate species [69–71]. Some of this activity is due to a mutation in the promotor region of the classical inducible form of *hsp70* resulting in constitutive expression and cold-enhanced activity of the proteosome in Antarctic notothenioids [72,73]. In our analyses on genes that were found to be under diversifying selection, we identified genes involved in protein folding and degradation such as members of heat shock protein family member (*hspa4*), ubiquilin/ubiquitin proteins (*ubqln4* and *ube2z*) and the chaperon *psmc1*. Similar to genes with acquired CNEs, such as the ubiquitin *ube2h*, *ube2e1*, *itch* and the co-chaperon *ahsa1*. Furthermore, there was evidence of selection pressures on genes involved in transcription, splicing, and the proteasome (e.g. *eif3b, sf3a2* and *psmb10*) indicating wider scale cold adaptation of the proteostasis pathway.

It should be noted that these examples are likely minimal identifications due to the stringency of orthologue assignment, which may not accurately identify duplicated genes, the number of which are considerable in the notothenioids [66]. Such analyses will also fail to identify those genes which are poorly conservated between species, such as genes involved in the immune function, intrinsically disordered proteins and those containing intrinsically disordered regions.

## Conclusions

This haplotype-resolved genome assembly of *H. antarcticus* thus provides a valuable resource for future functional investigations into cold adaptation and resilience to climate change. Overall haplotype diversity was low and contrary to expectations haplotypic diversity in the *afgp* locus was also low despite a high number of transposon insertions. However, we did identify sequence length variation in the *afgp* genes between the two haplotypes, which could be biologically relevant. Furthermore, we show that genomic gain was linked to transposon activity which correlates with drops in temperature in the Southern Ocean, and this activity resulted in the generation of novel genomic elements. Finally, we show that there has been significant selection pressure on notothenioid genomes, both in genes and control regions, to enable these fish to thrive for much of their life below 0°C.

## Methods

### Sequencing data generation

The specimen of *Harpagifer antarcticus* that was used for genome assembly was collected at Ryder Bay in the Antarctic peninsula (coordinates: −67,59053 −68,28863), and was preserved by flash freezing. This specimen was the same that was used for the generation of the fHarAnt1.1 assembly (accession: GCA_902827135.1; BioSample: SAMEA104132831) [6].

High Molecular Weight (HMW) DNA was extracted using the Sanger Tree of Life Manual MagAttract v1 protocol [74] in the Tree of Life Core Laboratory at the Wellcome Sanger Institute. To assess the quantity of the extracted DNA the Qubit^TM^ 1X dsDNA High Sensitivity assay (ThermoFisher, Cat No. Q33231) on the Qubit Flex Fluorometer (ThermoFisher, Cat No. Q33327) was used. The quality of the DNA was then determined using the Femto Pulse (Agilent, PN: M5330AA) instrument with the gDNA 165 kb Analysis kit 275 samples (Agilent, Cat No. FP-1002-0275). Approximately 3µg of the extracted DNA underwent library preparation for ONT sequencing using the Ligation Sequencing Kit V14 (SQK-LSK114). Sequencing was performed on one flow cell on the PromethION 24 in Sequencing Operations at the Wellcome Sanger Institute.

Three *H. antarcticus* specimens were Schedule 1 killed and the gills rapidly dissected, added to tissue culture media (L15 without phenol red, 10% foetal bovine serum and antibiotics) and placed on ice. The gills were chopped up in culture media and the resultant cell suspensions of each sample added to individual wells in 6-well Nunclon™ tissue culture plates. For each fish, one of each of the gill samples was incubated at 0°C with the paired sample incubated at 6°C for 6 days. Cells were harvested and RNA extracted using TRI Reagent according to manufacturer’s instructions. 1µg of rRNA depleted RNA was used for RNAseq library preps with Kapa Hyperprep UDI kit. (Cat#KK8544). Paired end sequencing was carried out by Novogene on the Novaseq PE150 platform.

### Genome assembly

The ONT reads were sequenced at an estimated 23.4X coverage per haplotype and based called with dorado 7.2.13 (https://github.com/nanoporetech/dorado) in super accurate mode. The reads were assembled using Hifiasm version 0.24.0-r702 [75] with --ont option in Hi-C phasing mode. This produced a phased assembly with contig N50 of 19.3 Mb and 14.8 Mb in each haplotype. The sizes of both haplotypes were 1.072 Gb and 1.222 Gb which were close to the kmer-based estimate from the GenomeScope at 1,152 Gb.

Furthermore Hi-C reads were mapped to each haplotype independently using bwa-mem2 [76]. Each haplotype was then scaffolded with Hi-C data in YaHS [77], using the --break option for handling potential mis-assemblies in contigs. The produced assembly was evaluated for kmer completeness using MerquryFK (https://github.com/thegenemyers/MERQURY.FK) using kmers from the ONT reads. This indicated 99.39% kmer completeness and QV value 42.6.

The individual haplotype assemblies were screened for contamination using the ASCC pipeline (https://github.com/sanger-tol/ascc) removing 5.8Mb Hap1 and 1.8Mb Hap2 non-target and mitochondrial sequences. The assemblies were then combined and prepared for curation using the treeval pipeline (https://github.com/sanger-tol/treeval) generating a Hi-C contact map with supplementary analysis tracks (telomeres, gaps and long read coverage). Assembly errors were corrected by manipulating the contact map in PretextView (https://github.com/sanger-tol/PretextView) an AGP of the resolved chromosomal haplotypes were exported and a curated fasta for each haplotype generated using pretext-to-asm (https://github.com/sanger-tol/agp-tpf-utils) with each super scaffold named according to descending size. Contact maps for each resolved haplotype were generated by curationpretext (https://github.com/sanger-tol/curationpretext) and images generated with PretextSnapshot (https://github.com/sanger-tol/PretextSnapshot).

### Repeat annotation

Repetitive elements were *de-novo* mined with RepeatModeler v.2.0.5 [78] adding the *-LTRStruct* extension to improve LTR detection. Potential multicopy host genes included in the repetitive library were removed with ProtExcluder v.1.2 after blasting (Blastx, e-value 1e-10) the repetitive library against a protein database consisting of all vertebrate proteins included in the Swiss-Prot database (The UniProt Consortium, 2025) and the predicted proteomes of five published notothenioid genomes available on NCBI RefSeq: *C. gobio, P.georgianus*, *Gymnodraco acuticeps, Trematomus bernacchii*, *Eleginops maclovinus*. Furthermore, a TE library was made with MCHelper v.1.7.0 with default parameters. Both haplotypes were annotated with RepeatMasker in sensitive mode (*-s*). RepeatMasker derived annotation was post-processed with: (a) parseRM.pl script (https://github.com/4ureliek/Parsing-RepeatMasker-Outputs/tree/master) to summarize the TE content and obtain TE landscapes describing transposon activity through relative time, and (b) the RM2Bed.py script from RepeatMasker to resolve overlaps in the TE annotation (--overlap resolution ‘higher score’) and produce bed files describing the coordinates of TE insertions.

Tandem repeats were identified with the Centrominer function of quarTeT v.1.2.5 [79]. rDNA genes were annotated with barrnap v.0.9 [80] https://github.com/tseemann/barrnap and telomeric repeats with tidk v0.2.31 [81] screening the assembled chromosomes for the TTAGGG telomeric motif.

### Gene annotation

Gene annotation of Hap1 was performed with BRAKER3 [82]. Repetitive elements were soft masked and both protein and transcriptomic data were used as external evidence to train AUGUSTUS [83] and GeneMark-ETP [84] *ab-initio* gene predictors. Illumina RNAseq reads from muscle, skin, liver, kidney, heart, and brain [5] (**Table S14**) and from gills (data generated herein) were mapped with HISAT2 [85]. The sorted bam files were supplied to BRAKER3 together with the protein database previously used for cleaning the repetitive libraries from host genes. The quality of the annotation was assessed with BUSCO v.5 [86]. To obtain a gene annotation for Hap2, we mapped BRAKER-derived genes with Liftoff [87] (-polish -copies) to obtain their lifted coordinates. Functional annotation was performed on the Hap1 predicted proteome with eggNOG-mapper v2 [88] transferring only annotations with experimental evidence and auto-adjusting the taxonomic scope based on the query sequence.

### Annotation of the *afgp* locus

To annotate the *afgp* locus on each haplotype, a combination of automated and manual approaches was used. The annotation of the *D. mawsoni afgp* locus (accession HQ447059.1, haplotype 1) [8] was used as a reference for homology-based gene prediction with GeMoMa v.1.7.1 [89] under default parameters. This initial annotation was supplemented by manually blasting *D. mawsoni* gene sequences with BLASTN v.2.12.0 [90] and reconstructing the gene structure based on alignment coordinates. Each identified *afgp* copy was then manually inspected for frameshifts and gaps to identify complete and pseudogenised gene copies. Finally, we used Klumpy [91] with the *find_klumps* function to verify the completeness of the annotation, using the exon-1 and exon-2 of one randomly chosen functional *afgp* gene copy previously annotated as query.

To infer evolutionary relationships of *afgp* genes, we used the short exon-1 (signal peptide) and intron-1 nucleotide sequences, excluding the highly repetitive exon-2s encoding the AFGP polyprotein, following the Nicodemus-Johnson et al., (2011) approach. Analysis was performed both with and without the chimeric *afgp/tlp* genes. Sequences were aligned with MAFFT v.7.525 [92] using the E-INS-i. Because *afgp* and *afgp/tlp* genes share only exon-1 and an initial part of intron-1, when these latest genes were included in the analyses, we removed the non-homologue region based on visual inspection of the alignment. Phylogenetic inference was performed with IQ-TREE 2 [93] with 1,000 ultrafast bootstrap replicates [94], to assess nodal support, selecting the best-fit evolutionary model with ModelFinder [95].

### Characterization of LINE-L2 elements

LINE-L2 insertions annotated on Hap1 and longer than 1,000 bp were extracted with bedtools *getfasta* and subjected to Blastx (-evalue 1e-05 –max_target_seqs 1 –max_hsps 1) against all reverse transcriptase (RT) amino acid (aa) sequences that compose the seed alignment of the Reverse transcriptase PFAM [96] profile PF00078. Based on the BLASTX v.2.12.0 [90] alignment coordinates, the amino acid sequences of the RT segments were extracted, discarding those shorter than 100 amino acids. To decrease the size of the dataset for the subsequent phylogenetic analyses all RT segments were clustered at 80% sequence identity with CD-HIT [97] keeping the longest element as representative of the cluster. All representative RT segments together with reference sequences of the LINE-L2, L2A, L2b, Crack, Daphne and CR1 clades described in [25] were aligned with MAFFT in G-INS-i mode and trimmed with Trimal v.1.4.1[98] in *gappyout* mode. Phylogenetic inference was performed with IQ-TREE v2.4.0 [93] with 1,000 UltraFast bootstrap replicates to assess nodal support and selecting the best-fit evolutionary model with ModelFinder.

CD-HIT [97] clustering identified eight clusters, each comprising between 600 and 3,669 RTs, together accounting for 73% of all extracted RTs longer than 100 aa. The complete insertion corresponding to the representative element of each one of these clusters was used as query in a “Blast-Extend-Extract" analyses [99–101] to reconstruct a consensus sequence representative of the family. The curated consensi were used in an additional RepeatMasker analyses, as previously described. We finally looked for the identified and reconstructed LINE-L2 family across genomes (**Table S5**) through BLASTN searches requiring an alignment length of at least 500 nucleotides with at least 80% of identity to the query consensus sequence.

### Haplotype comparison

A whole genome alignment of the two haplotypes was performed with Minimap2 v.2.26 [102] and inferred SNPs, SVs (insertions, deletions, and duplications), and rearranged genomic regions (inversions and translocations) between the two haplotypes using Syri v.1.7.1 [103]. In each case Hap1 was used as the reference. We identified genes and exons subject to structural variation between the two haplotypes by intersecting the gene annotation with Syri-derived SV and rearrangement coordinates using bedtools v2.30 [104] *intersect.* We tested whether the genomic distribution of SVs, considering only translocations, duplications and inversions, was associated with specific genomic features using a generalized linear mixed model (GLMM) with a binomial error structure. SVs were compared to an equal number of randomized genomic intervals generated with bedtools *shuffle*. For each interval we quantified the following: distance to the nearest chromosome end, distance to the closest assembly gap, length of the SV, and the proportion of bases within 50 kb flanking windows annotated as TEs, tandem repeats, or LINE elements. All predictors were standardized prior to analysis. The model included chromosome as a random intercept to account for non-independence of intervals on the same chromosome. GLMMs were fitted in R using the lme4 package [105]. Odds ratios and 95% confidence intervals were obtained by exponentiating model coefficients.

To identify putative heterozygous TE insertions, we overlapped the RepeatMasker annotations of Hap1 and Hap2 with deletions and insertion events, respectively, requiring that both insertions/deletions and the TE annotation overlap by at least 75% of their length.

To assess whether the SVs identified between haplotypes might result from assembly errors, we additionally mapped ONT reads back to the assembly with Minimap2 and called heterozygous SVs with Sniffles2 [106]. We then quantified the overlap between the genome-based and read-based heterozygous SVs larger than 50 bp using SURVIVOR [107]. Overlapping SVs genotyped as homozygous for the alternate allele were considered as putative mis-assemblies.

### Species tree reconstruction and divergence time estimation

Divergence times were estimated across a set of genomes of notothenioids, including our newly generated *H. antarcticus* assembly (Hap1), and 11 publicly available chromosomal assemblies: *Cottoperca gobio* (*Cottoperca trigloides*) [23]*, Notothenia rossii* [108], *Pseudochaenichthys georgianus* [6], *Eleginops maclovinus* [109], *Dissostichus mawsoni* [110], *Dissostichus eleginoides, Chaenocephalus aceratus* [111], *Pagothenia borchgrevinki* (*Trematomus borchgrevinki*) [112], *Champsocephalus gunnari, Champsocephalus esox* [9], *Pogonophryne albipinna* [113] (**Table S5**). Additionally another seven marine temperate and tropical fishes’ representative of four, closely related orders (**Table S5**).

First, we manually constructed a topology of the included species following the [6] and [9] inferred phylogenetic relationships. Branch lengths were then estimated using the nucleotide sequences of orthologous single-copy BUSCO genes extracted with BUSCO v.5.7.1 [86] and the actinopterygii_odb10 reference dataset. Briefly, single-copy genes present in all analysed species, were separately aligned using MAFFT [92] with the E-INS-i algorithm. Each alignment was cleaned of ambiguous positions using TrimAl in automated mode, and sequences sharing only a reduced region with the others in the alignment were removed using the parameters -resoverlap 0.8 and -seqoverlap 75. Finally, only alignments with at least 500 positions and for which no sequences were excluded by TrimAl were retained. To reduce computational time, 1,000 alignments were randomly selected, concatenated, and subjected to constrained phylogenetic inference with IQ-TREE v2.4.0, selecting the best-fit evolutionary model with ModelFinder. The resulting tree was dated with LSD2 [114] under a fixed substitution rate using [6] estimates as priors of the following nodes:1) split between *Takifugu rubripes* and *Nelusetta ayraudi* from all other species: 94 MYA, 2) notothenioid emergence: 47 MYA, and 3) cryonotothenioid diversification: 10.7 MYA. A neutral substitution rate of notothenioids was obtained by re-running the same analyses on the notothenioid-only subtree and using 4-fold degenerate sites extracted from the BUSCO concatenated alignment. The resulting neutral rate of 3.2E-3 million year was used to translate % of CpG corrected Kimura Divergence of TE landscape profiles into absolute time.

### Selection analyses

Publicly available predicted proteomes of fish species included in divergence time estimation were clustered into orthogroups (OG) with OrthoFinder v2.5.4 [115] in *ultra-sensitive* mode. We then isolated OGs in which at least one gene for all species was present. Multi-copy OGs (i.e. more than one gene was present for at least one species) were decomposed into single-copy OGs using DISCO [116]. Decomposed and original single-copy OGs were then aligned with Prank v250331 [117] under a codon model, cleaned of ambiguous positions with Gblocks taking into consideration the codon structure [118] and from sequences with high percentage of gappy positions with TrimAl (-resoverlap 0.6 -seqoverlap 55) [98]. After filtering, only OGs for which all species were present were kept (i.e. not affected by Trimal filtering) and with an alignment longer than 300 codon positions. IQ-TREE2 v2.4.0 was used to infer gene trees under a GTR+I+G substitution model. Because our aim was to detect genes that might be strictly related to cold adaptation in extant cryonotothenioids, we used aBSREL from the HyPhy package v2.5.8 [119] testing for diversifying selection acting on their stem branch. Prior to this, to ensure that we were testing the same evolutionary event we discarded any gene trees in which cryonotothenioids were not monophyletic and not in a sister relationship with their true temperate notothenioid sister species *E. maclovinus* using a custom R script. Finally, because aBSREL analyses does not test that selection has not occurred outside tested branches we re-ran it for all genes previously identified as under diversifying selection along the cryonotothenioid stem tagging all temperate-adapted species (including non-cryonotothenioid notothenioids) and all their ancestral branches. OGs that were found to be under positive selection in at least one temperate-adapted branch were then removed from aBSREL significant results on the cryonotothenioid ancestor. GO term enrichment was performed using all genes analysed by aBSREL as background based on EGG-NOG annotation with the TopGO R package [120] with a Fisher exact test and an *elim* algorithm.

### Synteny detection and multi-genome alignment of notothenioids

The same set of notothenioid genomes included in divergence time estimation was used to infer syntenic relationships based on gene order conservation using GENESPACE v1.2.3 [121] with default parameters and Progressive Cactus v2.9.7 [122] to perform a multi whole-genome alignment. Prior to running Progressive Cactus, the genome assemblies were soft masked with RepeatMasker, using a repeat library generated for each genome using RepeatModeler2 with default parameters. To guide the whole-genome alignment we used the notothenioid-only subtree obtained with the constrained tree search and the 1,000 randomly selected BUSCO genes.

We inferred insertion and deletion events along all branches of the species tree with the halBranchMutations function of the HAL package [123]. Net genome gain rates per MY were calculated by subtracting the total number of base pairs involved in deletion events from those involved in insertion events and dividing the result by the branch length in time. Insertions inferred along the *H. antarcticus* ancestors were then lifted off to the *H. antarcticus* Hap1 assembly using halLiftover (--noDupes). Due to the fragmentation of the lifted insertions, we merged intervals closer than 100 bp when they derive from the same ancestral insertion events. To detect putative TE-derived insertion events, lifted annotations were intersected with the MCHelper-derived TE annotation requiring a reciprocal overlap of the TE annotation and lifted insertion of 75%. PhyloFit from the PHAST package v1.5 [124] was used under a REV substitution model to train an initial non-conserved model of evolution on the same set of 4-fold degenerate sites extracted from BUSCO genes that was used to estimate the neutral substitution rate. PhastCons was then applied to a MAF representation of the whole-genome multiple alignment, generated with cactus-hal2maf using *H. antarcticus* as the reference genome (--noAncestors --refGenome HarAnt1.hap1 --dupeMode single), to refine the non-conserved model and train a conserved model for each chromosome. Parameters from the per-chromosome conserved and non-conserved models were then averaged using PhyloBoot, and PhastCons was re-run to predict discrete Conserved Elements (CEs). CEs closer than 5 bp were then merged into single elements. To identify CEs shared by all analysed notothenioids, and those that emerged along the cryonotothenioid stem branch, we used halAlignmentDepth to calculate per-base alignment depth for the *H. antarcticus* genome. Elements covered over at least 75% of their length by all other notothenioids were considered ancestrally present, whereas elements covered over at least 75% of their length only by cryonotothenioids were considered to have been acquired along their stem branch. We assigned each CE to different genomic features (exons, introns, promoters, transcription terminating site, and intergenic regions) with the HOMER annotatePeaks.pl script [125]. GO term enrichment analyses was performed similarly to what was performed for selection analyses but using all *H. antarcticus* genes as background.

## Supplementary Material

Supplementary Figures

Supplementary Tables

## Data availability

The datasets generated in the present study have been submitted on NCBI and will become public upon acceptance for publication. Biosample: SAMEA104132831.

## Acknowledgments

We thank the Wellcome Sanger Institute DNA pipelines for support with sequencing data generation of the *H. antarcticus* genome. We would like to thank the Rothera Marine Team for sample collection. IB and JM were supported by the Centre for Translational Biodiversity Genomics (LOEWE-TBG) funded through the program LOEWE–Landes-Offensive zur Entwicklung Wissenschaftlich-ökonomischer Exzellenz of Hesse’s Ministry of Higher Education, Research, and the Arts. NF and DB were supported by NIH grant R35 GM144336.

## Author Contributions

I.B. conceptualised the study and supervised the work. I.B., J.M. designed analysis. A.D., M.S.C., N.F., I.B., generated data. I.B., R.D., D.L.B., N.F. contributed resources, K.K. performed genome assembly, T.M., J.M.D.W. performed genome curation, J.M. performed data analysis, with J.M.D.W input. J.M. and I.B. wrote the original manuscript with contributions from M.S.C. All co-authors read, provided feedback, and approved the manuscript.

