## Supplementary_Information for "A new phased assembly of the Antarctic spiny plunderfish provides novel insights into the evolution of the notothenioid radiation"

Supplementary Tables

**Supplementary Table 1**

| | **Hap1** | **Hap2** |
|---|---|---|
| **Total length** | 1.068.962.898 | 1.220.510.434 |
| **Scaff number** | 1.435 | 653 |
| **Contig N50** | 17.034.000 | 12.766.000 |
| **Scaf N50** | 46.795.402 | 42.703.524 |
| **Number of Chromosomes** | 24 | 24 |
| **BUSCO** | 99.0%[S:98.4%,D:0.6%],F:0.6%,M:0.4%,n:3640 | 99.0%[S:98.3%,D:0.7%],F:0.7%,M:0.3%,n:3640 |
| **QV** | 40 | 40,4 |

**Supplementary Table 2**

| **Scaffold** | **Chromosome** |
|---|---|
| SUPER_4 | 1 |
| SUPER_24 | 2 |
| SUPER_2 | 3 |
| SUPER_17 | 4 |
| SUPER_12 | 5 |
| SUPER_8 | 6 |
| SUPER_3 | 7 |
| SUPER_13 | 8 |
| SUPER_1 | 9 |
| SUPER_9 | 10 |
| SUPER_16 | 11 |
| SUPER_20 | 12 |
| SUPER_6 | 13 |
| SUPER_18 | 14 |
| SUPER_7 | 15 |
| SUPER_5 | 16 |
| SUPER_10 | 17 |
| SUPER_23 | 18 |
| SUPER_21 | 19 |
| SUPER_22 | 20 |
| SUPER_11 | 21 |
| SUPER_14 | 22 |
| SUPER_19 | 23 |
| SUPER_15 | 24 |

**Supplementary Table 3**

| | Hap1 | Hap2 |
|---|---|---|
| **Number of genes** | 23,057 | 22,463 |
| **Number of mRNAs** | 48,897 | 31,527 |
| **Mean number of introns per mRNA** | 8.3 | 8.4 |
| **Mean number of exons per mRNA** | 9.3 | 9.4 |
| **Mean gene length** | 14,904 | 14,715 |
| **Mean intron length** | 1,555 | 1,522 |
| **Mean exon length** | 174 | 124 |
| **BUSCO (actinopterygii odb_10)** | C:98.9%[S:97.7%,D:1.2%],F:0.5%,M:0.6% | C:96.7%[S:95.6%,D:1.1%],F:1.1%,M:2.2% |

**Supplementary Table 4**

| Overview table | | |
|---|---|---|
| **TE class** | **Hap1** | **Hap2** |
| **LINEs** | 13 | 11,34 |
| **LTRs** | 13,98 | 17,84 |
| **DNA** | 13,88 | 12,25 |
| **RC** | 0,05 | 0,04 |
| **SINE** | 0,06 | 0,05 |
| **Unknown** | 0,2 | 0,18 |
| **Tandem repeats** | 4,5 | 4,8 |
| **Simple repeats** | 2,35 | 2,07 |
| **Low complexity** | 0,15 | 0,14 |
| **Total** | 43,67 | 43,91 |

| TEs by Superfamily | Genome coverage (%) | | |
|---|---|---|---|
| **Superfamily** | **Hap1** | **Hap2** | **Class** |
| **TcMar-ISRm11** | 0,004779691 | 0,004406599 | DNA |
| **Sola-1** | 0,005902255 | 0,005421994 | DNA |
| **MULE-NOF** | 0,006541181 | 0,006372416 | DNA |
| **R2-NeSL** | 0,009075744 | 0,008463754 | LINE |
| **hAT-hATx** | 0,010835457 | 0,00946604 | DNA |
| **I-Jockey** | 0,012170373 | 0,010227934 | LINE |
| **RTE-X** | 0,013299111 | 0,013012916 | LINE |
| **TcMar-Fot1** | 0,01331492 | 0,011887567 | DNA |
| **hAT-Tag1** | 0,016618907 | 0,014290496 | DNA |
| **TcMar-Tc2** | 0,01901053 | 0,015485161 | DNA |
| **CR1** | 0,019605302 | 0,017100141 | LINE |
| **hAT-hAT5** | 0,020206248 | 0,017028122 | DNA |
| **Zisupton** | 0,030205956 | 0,026711119 | DNA |
| **Penelope** | 0,043723406 | 0,041617096 | Penelope |

| | | | |
|---|---|---|---|
| **Helitron** | 0,045550098 | 0,041057248 | RC |
| **Dada** | 0,04571287 | 0,062060018 | DNA |
| **PIF-ISL2EU** | 0,053272965 | 0,046735528 | DNA |
| **Unknown** | 0,05527562 | 0,046773218 | SINE |
| **Merlin** | 0,055644008 | 0,04861294 | DNA |
| **Proto2** | 0,056050657 | 0,047631055 | LINE |
| **Crypton-H** | 0,065469906 | 0,05653356 | DNA |
| **Ginger-1** | 0,077340741 | 0,06877434 | DNA |
| **Ngaro** | 0,079720203 | 0,071085996 | LTR |
| **Crypton-A** | 0,088138124 | 0,076345271 | DNA |
| **PIF-Harbinger** | 0,097676365 | 0,077930264 | DNA |
| **Copia** | 0,098900895 | 0,085675957 | LTR |
| **Sola-2** | 0,099819246 | 0,086373289 | DNA |
| **IS3EU** | 0,126678184 | 0,110161533 | DNA |
| **hAT-Blackjack** | 0,128556888 | 0,113959288 | DNA |
| **Kolobok-E** | 0,147513816 | 0,319307471 | DNA |
| **CMC-EnSpm** | 0,148790452 | 0,136953848 | DNA |
| **ERV** | 0,15933582 | 0,141425665 | LTR |
| **Unknown** | 0,20210991 | 0,177821585 | Unknown |
| **Maverick** | 0,239393727 | 0,207085079 | DNA |
| **P** | 0,250950338 | 0,218685472 | DNA |
| **Unknown** | 0,326903964 | 0,27877443 | LINE |
| **L1-Tx1** | 0,337941388 | 0,26045185 | LINE |
| **L1** | 0,343850379 | 0,316897578 | LINE |
| **Pao** | 0,346597293 | 0,293703266 | LTR |
| **Kolobok-T2** | 0,364417532 | 0,319307471 | DNA |
| **PiggyBac** | 0,51869938 | 0,448819842 | DNA |
| **hAT-Tip100** | 0,554194204 | 0,484122203 | DNA |
| **RTE-BovB** | 0,612075295 | 0,542266565 | LINE |
| **DIRS** | 0,660656504 | 0,579024546 | LTR |
| **I** | 0,684329604 | 0,577757126 | LINE |
| **hAT-Charlie** | 0,685612134 | 0,610916039 | DNA |
| **TcMar-Tc1** | 0,751543413 | 0,662926983 | DNA |
| **Rex-Babar** | 0,919560583 | 0,788629637 | LINE |
| **hAT-Ac** | 1,029050438 | 0,903821606 | DNA |

| | | | |
|---|---|---|---|
| **ERV1** | 1,093929033 | 0,957094071 | LTR |
| **Gypsy** | 1,999559437 | 1,72982667 | LTR |
| **Unknown** | 8,221667769 | 7,274402621 | DNA |
| **Unknown** | 9,539620475 | 13,98632656 | LTR |
| **L2** | 9,630117667 | 8,437966619 | LINE |

### Supplementary Table 5a

| Species | NCBI taxon ID | Assembly ID | Family | size (Mb) | Assembly size | Contig N50 (Mb) | Scaffold N50 (Mb) | BUSCO | Assembly Accession | Reference |
|---|---|---|---|---|---|---|---|---|---|---|
| *Cottoperca gobio (Cottoperca trigloides)* | 56716 | fCotGob3.1 | Bovichtidae | 609 | 609.4 | 6,3 | 25,2 | C:93.9%[S:92.5%,D:1.4%],F:0.9%,M:5.2% | GCF_900634415.1 | Bista et al. 2020 |
| *Eleginops maclovinus* | 56733 | JC_Emac_rtc_rv5 | Eleginopsidae | 606 | 606.3 | 7,6 | 26,7 | C:98.0%[S:97.0%,D:1.0%],F:0.4%,M:1.6% | GCF_036324505.1 | Cheng et al. 2024 |
| *Dissostichus mawsonii* | 36200 | KU_Dm_1.0 | Nototheniidae / Pleuragrammatinae | 926 | 926.4 | 3 | 37 | C:95.6%[S:92.1%,D:3.5%],F:1.7%,M:2.7% | GCA_011823955.1 | Lee at al., 2021 |
| *Dissostichus eleginoides* | 100907 | KU_De_1.0 | Nototheniidae / Pleuragrammatinae | 843 | 844.7 | 4,2 | 36 | C:97.8%[S:94.7%,D:3.1%],F:1.0%,M:1.2% | GCA_031216635.1 | Lee et al. 2023 |
| *Pagothenia borchgrevinki (Trematomus borchgrevinki)* | 8213 | PborFfbFv9 | Nototheniidae / Trematominae | 935 | 935.1 | 1,8 | 42,7 | C:98.0%[S:96.4%,D:1.6%],F:0.7%,M:1.3% | GCA_044885175.1 | Rayamajhi et al. 2025 |
| *Notothenia rossii* | 101497 | fNotRos5.1 | Nototheniidae / Nototheniinae | 1000 | 1,043 | 0,383 | 89,7 | C:94.8%[S:94.0%,D:0.8%],F:2.0%,M:3.2% | GCA_949606895.1 | Bista et al. 2024 |
| *Harpagifer antarcticus* | 43256 | fHarAnt1.2 | Harpagiferidae | 1000 | 1.072 / 1.222 | 17 | 44,8 | C:99.0%[S:98.4%,D:0.6%],F:0.6%,M:0.4% | tbc | this manuscript |
| *Pogonophryne albipinna* | 1090488 | KU_S6 | Artedidraconidae | 1100 | 1.074 | 0,962 | 41,8 | C:97.0%[S:94.3%,D:2.7%],F:0.9%,M:2.1% | GCA_028583405.1 | Jo et al. 2023 |
| *Pseudochaenichthys georgianus* | 52239 | fPseGeo1.2 | Channichthyidae | 1100 | 1.026 | 0,657 | 42,8 | C:94.1%[S:93.0%,D:1.1%],F:1.3%,M:4.6% | GCF_902827115.2 | Bista et al., 2023 |

### Supplementary Table 5b

| Species | NCBI taxon ID | Assembly ID | Family | size (Mb) | Assembly size | Contig N50 (Mb) | Scaffold N50 (Mb) | BUSCO | Assembly Accession | Reference |
|---|---|---|---|---|---|---|---|---|---|---|
| *Champsocephalus gunnari* | 52237 | JC_Cgun_ftc_fv8 | Channichthyidae | 994 | 994.2 | 3,2 | 44,1 | C:97.8%[S:96.2%,D:1.6%],F:0.7%,M:1.5% | GCA_036324595.1 | River-Colon et al., 2023 |
| *Champsocephalus esox* | 159716 | JC_Ceso_ftc_fv8 | Channichthyidae | 987 | 987.1 | 2,6 | 43,6 | C:97.3%[S:96.2%,D:1.1%],F:1.0%,M:1.7% | GCA_036324585.1 | River-Colon et al., 2023 |
| *Chaenocephalus aceratus* | 36190 | KU_Ca_2.0 | Channichthyidae | 1100 | 1.065 | 1,5 | 33,5 | C:92.6%[S:88.9%,D:3.7%],F:1.5%,M:5.9% | GCA_023974075.1 | Lee et al., 2023 |
| *Gasterosteus aculeatus* | 481459 | GAculeatus_UGA_version5 | Gasterosteidae | 472 | 471.9 | 0.4858 | 20.4 | C:97.0%[S:94.6%,D:2.4%],F:1.1%,M:1.9% | GCF_016920845.1 | Nath et al., 2021 |
| *Pungitius pungitius* | 134920 | fPunPun2.1 | Gasterosteidae | 480 | 480.4 | 1.4 | 21 | C:97.8%[S:97.0%,D:0.8%],F:0.8%,M:1.4% | GCF_949316345.1 | Hänfling et al., 2023 |
| *Larimichthys crocea* | 215358 | L_crocea_2.0 | Sciaenidae | 658 | 657.9 | 0.2775 | 27 | C:98.4%[S:97.5%,D:0.9%],F:0.5%,M:1.1% | GCF_000972845.2 | Ao et al., 2015 |
| *Labrus bergylta* | 56723 | fLabBer1.1 | Labridae | 720 | 720.2 | 2.7 | 31.3 | C:97.8%[S:97.1%,D:0.7%],F:0.6%,M:1.6% | GCF_963930695.1 | NA |
| *Notolabrus celidotus* | 1203425 | fNotCel1.pri | Labridae | 847 | 846.7 | 3.7 | 37.1 | C:95.8%[S:94.9%,D:0.9%],F:0.6%,M:3.6% | GCF_009762535.1 | NA |
| *Nelusetta ayraudi* | 303726 | CSIRO-AGI_Nayr_v1 | Monacanthidae | 577 | 576.5 | 6.9 | 6.9 | C:96.7%[S:95.7%,D:1.0%],F:1.0%,M:2.3% | GCF_046127955.1 | NA |
| *Takifugu rubripes* | 31033 | fTakRub1.2 | Tetraodontidae | 384 | 384.1 | 3.1 | 16.7 | C:96.9%[S:94.3%,D:2.6%],F:0.7%,M:2.4% | GCF_901000725.2 | NA |

**Supplementary Table 6**

| Variant type | Number of events | Length reference | Length query |
|---|---|---|---|
| **Inversions** | 155 | 8.868.241 | 9.134.047 |
| **Translocations** | 145 | 4.197.546 | 3.647.854 |
| **Duplications** | 251 | 4.609.203 | - |
| **Insertions** | 214.681 | | 16.640.238 |
| **Deletions** | 214.893 | 16.808.012 | - |
| **SNPs** | 1.620.216 | 52.762.169 | 50.471.908 |
| **Highly diverged regions** | 24.540 | 14.694 | 14.694 |

**Supplementary Table 7**

| | Split | Deletions (bp) | Insertions (bp) | Net gain (Mb) | Branch length (MY) | Net gain rate (Mb X MY) |
|---|---|---|---|---|---|---|
| **Anc01** | EleMac - cryonotothenioids | 499043 | 100799664 | 100,30 | 20.7 | 4.84544062801932 |
| **Anc02** | Cryonotothenioid ancestor | 5942195 | 262680582 | 256,74 | 15.6 | 16.4575889102564 |
| **Anc03** | DisMaw - DisEle | 1915618 | 98933047 | 97,02 | 4.65 | 20.8639632258065 |
| **Anc04** | PagBor - NotRos | 944905 | 20853349 | 19,91 | 1.2 | 16.59037 |
| **Anc05** | NotRos - PogAlb | 7118060 | 22274394 | 15,16 | 1.48 | 10.2407662162162 |
| **Anc06** | PogAlb - ChaGun | 7995185 | 59483192 | 51,49 | 1.87 | 27.5336935828877 |
| **Anc07** | PogAlb - HarAnt | 3246574 | 53989080 | 50,74 | 1.2 | 42.2854216666667 |
| **Anc08** | ChaGun - PseGeo | 6099811 | 111093996 | 104,99 | 2.12 | 49.5255589622641 |
| **Anc09** | ChaGun - ChaEso | 4932407 | 163657480 | 158,73 | 2.09 | 75.9450110047847 |
| **Anc10** | PseGeo - ChaAce | 2868300 | 62970793 | 60,10 | 0.79 | 76.0791050632911 |
| **ChaAce** | / | 5511882 | 147085816 | 141,57 | 3.24 | 43.6956586419753 |
| **ChaEso** | / | 6638005 | 106860420 | 100,22 | 1.93 | 51.9287124352332 |
| **ChaGun** | / | 3755646 | 99118512 | 95,36 | 1.93 | 49.4108113989637 |
| **CotGob** | / | 5227 | 187530812 | 187,53 | 47 | 3.98990606382979 |
| **DisEle** | / | 4580041 | 87105609 | 82,53 | 6.05 | 13,6405897520661 |
| **DisMaw** | / | 4639833 | 132781946 | 128,14 | 6.05 | 21,1805145454545 |
| **EleMac** | / | 8496818 | 139792732 | 131,30 | 26.3 | 4.99224007604563 |
| **HarAnt** | / | 10392579 | 351876096 | 341,48 | 4.95 | 68.9865690909091 |
| **NotRos** | / | 10414889 | 399848994 | 389,43 | 8.01 | 48.6184900124844 |
| **PagBor** | / | 13108365 | 278777704 | 265,67 | 9.5 | 27.9651935789474 |
| **PogAlb** | / | 5204039 | 240104298 | 234,90 | 4.95 | 47.4545977777778 |
| **PseGeo** | / | 5467138 | 217171889 | 211,70 | 3.24 | 65.3409725308642 |

**Supplementary Table 8**

| Node | Split | DNA | LINE | LTR | Total | Branch length | Rate x MY |
|---|---|---|---|---|---|---|---|
| **Anc01** | EleMac - cryonotothenioids | 3342 | 1469 | 1173 | 5984 | 20,7 | 289,0821256 |
| **Anc02** | Cryonotothenioid ancestor | 9635 | 9767 | 3992 | 23394 | 15,6 | 1499,615385 |
| **Anc04** | PagBor - NotRos | 1579 | 991 | 537 | 3107 | 1,2 | 2589,166667 |
| **Anc05** | NotRos - PogAlb | 4320 | 3092 | 1315 | 8727 | 1,48 | 5896,621622 |
| **Anc06** | PogAlb - ChaGun | 8442 | 7314 | 2581 | 18337 | 1,87 | 9805,882353 |
| **Anc07** | PogAlb - HarAnt | 4040 | 2864 | 1339 | 8243 | 1,2 | 6869,166667 |
| **HarAnt** | / | 16994 | 23600 | 11163 | 51757 | 4,95 | 10455,9596 |

**Supplementary Table 9**

| GO.ID | Term | Annotated | Significant | Expected | classicFisher |
|---|---|---|---|---|---|
| GO:0060021 | roof of mouth development | 84 | 15 | 4,97 | 0,00017 |
| GO:0010761 | fibroblast migration | 38 | 8 | 2,25 | 0,00024 |
| GO:0035019 | somatic stem cell population maintenance | 56 | 11 | 3,31 | 0,00037 |
| GO:0008542 | visual learning | 56 | 11 | 3,31 | 0,00037 |
| GO:0045666 | positive regulation of neuron differentiation | 414 | 42 | 24,47 | 0,00042 |
| GO:0007399 | nervous system development | 2406 | 205 | 142,22 | 0,00046 |
| GO:0023052 | signaling | 5150 | 373 | 304,41 | 0,00049 |
| GO:0007267 | cell-cell signaling | 1389 | 115 | 82,1 | 0,00051 |
| GO:0072176 | nephric duct development | 19 | 6 | 1,12 | 0,00058 |
| GO:0001655 | urogenital system development | 381 | 41 | 22,52 | 0,0006 |
| GO:0045777 | positive regulation of blood pressure | 40 | 8 | 2,36 | 0,00079 |
| GO:0043401 | steroid hormone receptor signaling pathway | 180 | 19 | 10,64 | 0,00079 |
| GO:0030099 | myeloid cell differentiation | 407 | 31 | 24,06 | 0,00089 |
| GO:0006367 | transcription initiation at RNA polymerase II promoter | 203 | 22 | 12 | 0,00095 |
| GO:0042474 | middle ear morphogenesis | 21 | 6 | 1,24 | 0,00105 |
| GO:0009415 | response to water | 21 | 6 | 1,24 | 0,00105 |
| GO:0010628 | positive regulation of gene expression | 1909 | 140 | 112,84 | 0,00112 |
| GO:0050810 | regulation of steroid biosynthetic process | 82 | 9 | 4,85 | 0,0012 |
| GO:0051965 | positive regulation of synapse assembly | 64 | 11 | 3,78 | 0,0012 |
| GO:0048384 | retinoic acid receptor signaling pathway | 33 | 8 | 1,95 | 0,00128 |
| GO:0048865 | stem cell fate commitment | 15 | 5 | 0,89 | 0,0013 |
| GO:0045987 | positive regulation of smooth muscle contraction | 38 | 8 | 2,25 | 0,00144 |

| | | | | | |
|---|---|---|---|---|---|
| GO:0009952 | anterior/posterior pattern specification | 261 | 29 | 15,43 | 0,00168 |
| GO:0010718 | positive regulation of epithelial to mesenchymal transition | 39 | 8 | 2,31 | 0,00172 |
| GO:0033574 | response to testosterone | 67 | 11 | 3,96 | 0,00174 |
| GO:0035850 | epithelial cell differentiation involved in kidney development | 48 | 9 | 2,84 | 0,00175 |
| GO:0051354 | negative regulation of oxidoreductase activity | 23 | 6 | 1,36 | 0,00177 |
| GO:0050869 | negative regulation of B cell activation | 34 | 6 | 2,01 | 0,0018 |
| GO:0030326 | embryonic limb morphogenesis | 134 | 15 | 7,92 | 0,00182 |
| GO:0050769 | positive regulation of neurogenesis | 510 | 47 | 30,15 | 0,00186 |
| GO:1902305 | regulation of sodium ion transmembrane transport | 53 | 7 | 3,13 | 0,00187 |
| GO:0030278 | regulation of ossification | 196 | 24 | 11,59 | 0,00213 |
| GO:0009950 | dorsal/ventral axis specification | 24 | 6 | 1,42 | 0,00225 |
| GO:0048670 | regulation of collateral sprouting | 24 | 6 | 1,42 | 0,00225 |
| GO:0003016 | respiratory system process | 33 | 6 | 1,95 | 0,00243 |
| GO:0001764 | neuron migration | 173 | 22 | 10,23 | 0,00269 |
| GO:0050919 | negative chemotaxis | 42 | 8 | 2,48 | 0,00282 |
| GO:0050768 | negative regulation of neurogenesis | 311 | 33 | 18,38 | 0,00303 |
| GO:0048368 | lateral mesoderm development | 18 | 5 | 1,06 | 0,0032 |
| GO:0002063 | chondrocyte development | 34 | 7 | 2,01 | 0,00324 |
| GO:0002062 | chondrocyte differentiation | 108 | 16 | 6,38 | 0,00336 |
| GO:0060669 | embryonic placenta morphogenesis | 31 | 7 | 1,83 | 0,00346 |
| GO:0007187 | G protein-coupled receptor signaling pathway, coupled to cyclic nucleotide second messenger | 187 | 21 | 11,05 | 0,00347 |
| GO:0032330 | regulation of chondrocyte differentiation | 61 | 9 | 3,61 | 0,00405 |
| GO:0001706 | endoderm formation | 68 | 9 | 4,02 | 0,00406 |
| GO:0060914 | heart formation | 27 | 6 | 1,6 | 0,00425 |
| GO:0003151 | outflow tract morphogenesis | 81 | 13 | 4,79 | 0,00451 |
| GO:0010470 | regulation of gastrulation | 55 | 9 | 3,25 | 0,00461 |

| | | | | | |
|---|---|---|---|---|---|
| GO:0003413 | chondrocyte differentiation involved in endochondral bone morphogenesis | 20 | 5 | 1,18 | 0,00525 |
| GO:0030857 | negative regulation of epithelial cell differentiation | 37 | 7 | 2,19 | 0,00532 |
| GO:1902742 | apoptotic process involved in development | 42 | 7 | 2,48 | 0,00597 |
| GO:0001541 | ovarian follicle development | 89 | 12 | 5,26 | 0,00598 |
| GO:0030282 | bone mineralization | 98 | 14 | 5,79 | 0,00605 |
| GO:0048662 | negative regulation of smooth muscle cell proliferation | 45 | 7 | 2,66 | 0,00616 |
| GO:0021983 | pituitary gland development | 60 | 8 | 3,55 | 0,00621 |
| GO:0030540 | female genitalia development | 21 | 5 | 1,24 | 0,00656 |
| GO:0086012 | membrane depolarization during cardiac muscle cell action potential | 21 | 5 | 1,24 | 0,00656 |
| GO:0048745 | smooth muscle tissue development | 21 | 5 | 1,24 | 0,00656 |
| GO:0086091 | regulation of heart rate by cardiac conduction | 39 | 7 | 2,31 | 0,00718 |
| GO:2000772 | regulation of cellular senescence | 30 | 6 | 1,77 | 0,00732 |
| GO:1902930 | regulation of alcohol biosynthetic process | 69 | 7 | 4,08 | 0,00753 |
| GO:0033339 | pectoral fin development | 22 | 5 | 1,3 | 0,00808 |
| GO:0060706 | cell differentiation involved in embryonic placenta development | 22 | 5 | 1,3 | 0,00808 |
| GO:0001893 | maternal placenta development | 35 | 8 | 2,07 | 0,00909 |
| GO:0038179 | neurotrophin signaling pathway | 38 | 5 | 2,25 | 0,0092 |
| GO:0055012 | ventricular cardiac muscle cell differentiation | 23 | 4 | 1,36 | 0,0092 |
| GO:0060285 | cilium-dependent cell motility | 74 | 7 | 4,37 | 0,0092 |
| GO:0007156 | homophilic cell-cell adhesion | 72 | 10 | 4,26 | 0,00945 |
| GO:0009607 | response to biotic stimulus | 1035 | 57 | 61,18 | 0,00945 |
| GO:0035148 | tube formation | 177 | 20 | 10,46 | 0,0095 |
| GO:0071229 | cellular response to acid chemical | 210 | 19 | 12,41 | 0,00967 |
| GO:0060324 | face development | 63 | 10 | 3,72 | 0,00969 |
| GO:0072102 | glomerulus morphogenesis | 15 | 4 | 0,89 | 0,00979 |
| GO:0035357 | peroxisome proliferator activated receptor signaling pathway | 15 | 4 | 0,89 | 0,00979 |

| | | | | | |
|---|---|---|---|---|---|
| GO:0039019 | pronephric nephron development | 15 | 4 | 0,89 | 0,00979 |
| GO:0019371 | cyclooxygenase pathway | 15 | 4 | 0,89 | 0,00979 |
| GO:0045622 | regulation of T-helper cell differentiation | 23 | 5 | 1,36 | 0,00983 |
| GO:0072202 | cell differentiation involved in metanephros development | 23 | 5 | 1,36 | 0,00983 |
| GO:0019395 | fatty acid oxidation | 99 | 8 | 5,85 | 0,00983 |
| GO:0090189 | regulation of branching involved in ureteric bud morphogenesis | 25 | 5 | 1,48 | 0,01001 |
| GO:0061217 | regulation of mesonephros development | 28 | 5 | 1,66 | 0,01002 |
| GO:0014020 | primary neural tube formation | 102 | 9 | 6,03 | 0,01003 |
| GO:0002684 | positive regulation of immune system process | 764 | 46 | 45,16 | 0,01008 |
| GO:0000902 | cell morphogenesis | 1013 | 75 | 59,88 | 0,0102 |
| GO:0032879 | regulation of localization | 2370 | 145 | 140,09 | 0,01026 |
| GO:0043065 | positive regulation of apoptotic process | 604 | 45 | 35,7 | 0,01046 |
| GO:0098609 | cell-cell adhesion | 573 | 57 | 33,87 | 0,01062 |
| GO:2000027 | regulation of animal organ morphogenesis | 218 | 21 | 12,89 | 0,01102 |
| GO:0007398 | ectoderm development | 24 | 5 | 1,42 | 0,01183 |
| GO:0048557 | embryonic digestive tract morphogenesis | 24 | 5 | 1,42 | 0,01183 |
| GO:0030325 | adrenal gland development | 43 | 7 | 2,54 | 0,01226 |
| GO:0010765 | positive regulation of sodium ion transport | 38 | 5 | 2,25 | 0,01247 |
| GO:0031667 | response to nutrient levels | 602 | 39 | 35,58 | 0,01264 |
| GO:0035113 | embryonic appendage morphogenesis | 143 | 18 | 8,45 | 0,01287 |
| GO:0051896 | regulation of phosphatidylinositol 3-kinase/protein kinase B signal transduction | 190 | 16 | 11,23 | 0,0131 |
| GO:0043433 | negative regulation of DNA-binding transcription factor activity | 132 | 15 | 7,8 | 0,01312 |
| GO:0010092 | specification of animal organ identity | 32 | 5 | 1,89 | 0,01318 |
| GO:0090504 | epiboly | 46 | 4 | 2,72 | 0,01326 |

| | | | | | |
|---|---|---|---|---|---|
| GO:0001708 | cell fate specification | 112 | 13 | 6,62 | 0,01368 |
| GO:0071880 | adenylate cyclase-activating adrenergic receptor signaling pathway | 25 | 5 | 1,48 | 0,01408 |
| GO:0060045 | positive regulation of cardiac muscle cell proliferation | 25 | 5 | 1,48 | 0,01408 |
| GO:0045618 | positive regulation of keratinocyte differentiation | 17 | 4 | 1 | 0,01553 |
| GO:0060307 | regulation of ventricular cardiac muscle cell membrane repolarization | 17 | 4 | 1 | 0,01553 |
| GO:1905941 | positive regulation of gonad development | 17 | 4 | 1 | 0,01553 |
| GO:0072234 | metanephric nephron tubule development | 17 | 4 | 1 | 0,01553 |
| GO:0031952 | regulation of protein autophosphorylation | 50 | 5 | 2,96 | 0,0156 |
| GO:0030279 | negative regulation of ossification | 71 | 9 | 4,2 | 0,01644 |
| GO:0007190 | activation of adenylate cyclase activity | 26 | 5 | 1,54 | 0,01661 |
| GO:0060009 | Sertoli cell development | 26 | 5 | 1,54 | 0,01661 |
| GO:0043406 | positive regulation of MAP kinase activity | 280 | 29 | 16,55 | 0,01699 |
| GO:0048536 | spleen development | 46 | 7 | 2,72 | 0,0175 |
| GO:0032526 | response to retinoic acid | 149 | 18 | 8,81 | 0,01752 |
| GO:0030501 | positive regulation of bone mineralization | 36 | 6 | 2,13 | 0,01781 |
| GO:0019933 | cAMP-mediated signaling | 143 | 15 | 8,45 | 0,01795 |
| GO:0034105 | positive regulation of tissue remodeling | 30 | 5 | 1,77 | 0,018 |
| GO:0050673 | epithelial cell proliferation | 361 | 36 | 21,34 | 0,01806 |
| GO:0021537 | telencephalon development | 289 | 31 | 17,08 | 0,01835 |
| GO:0051347 | obsolete positive regulation of transferase activity | 602 | 53 | 35,58 | 0,01858 |
| GO:0045616 | regulation of keratinocyte differentiation | 35 | 8 | 2,07 | 0,01881 |
| GO:0014912 | negative regulation of smooth muscle cell migration | 18 | 4 | 1,06 | 0,01906 |

| | | | | | |
|---|---|---|---|---|---|
| GO:0061333 | renal tubule morphogenesis | 82 | 11 | 4,85 | 0,01911 |
| GO:0001657 | ureteric bud development | 96 | 12 | 5,67 | 0,01912 |
| GO:0002822 | regulation of adaptive immune response based on somatic recombination of immune receptors built from immunoglobulin superfamily domains | 110 | 9 | 6,5 | 0,01929 |
| GO:0090090 | negative regulation of canonical Wnt signaling pathway | 116 | 13 | 6,86 | 0,01933 |
| GO:0060444 | branching involved in mammary gland duct morphogenesis | 27 | 5 | 1,6 | 0,01942 |
| GO:0007413 | axonal fasciculation | 27 | 5 | 1,6 | 0,01942 |
| GO:0060992 | response to fungicide | 27 | 5 | 1,6 | 0,01942 |
| GO:0006482 | protein demethylation | 27 | 5 | 1,6 | 0,01942 |
| GO:0046697 | decidualization | 27 | 5 | 1,6 | 0,01942 |
| GO:0045665 | negative regulation of neuron differentiation | 246 | 23 | 14,54 | 0,01975 |
| GO:0048639 | positive regulation of developmental growth | 184 | 18 | 10,88 | 0,01983 |
| GO:0048169 | regulation of long-term neuronal synaptic plasticity | 37 | 6 | 2,19 | 0,02022 |
| GO:0001701 | in utero embryonic development | 372 | 38 | 21,99 | 0,02046 |
| GO:0006469 | negative regulation of protein kinase activity | 242 | 22 | 14,3 | 0,02057 |
| GO:0010976 | positive regulation of neuron projection development | 319 | 28 | 18,86 | 0,02204 |
| GO:0043392 | negative regulation of DNA binding | 50 | 7 | 2,96 | 0,02244 |
| GO:2000378 | negative regulation of reactive oxygen species metabolic process | 60 | 7 | 3,55 | 0,02251 |
| GO:0051968 | positive regulation of synaptic transmission, glutamatergic | 28 | 5 | 1,66 | 0,02252 |
| GO:0030890 | positive regulation of B cell proliferation | 28 | 5 | 1,66 | 0,02252 |
| GO:0086005 | ventricular cardiac muscle cell action potential | 28 | 5 | 1,66 | 0,02252 |
| GO:0021871 | forebrain regionalization | 28 | 5 | 1,66 | 0,02252 |
| GO:0021952 | central nervous system projection neuron axonogenesis | 28 | 5 | 1,66 | 0,02252 |

| | | | | | |
|---|---|---|---|---|---|
| GO:0002070 | epithelial cell maturation | 19 | 4 | 1,12 | 0,02304 |
| GO:0033334 | fin morphogenesis | 19 | 4 | 1,12 | 0,02304 |
| GO:0048608 | reproductive structure development | 490 | 54 | 28,96 | 0,02343 |
| GO:0035137 | hindlimb morphogenesis | 45 | 8 | 2,66 | 0,0235 |
| GO:0055074 | calcium ion homeostasis | 401 | 28 | 23,7 | 0,02358 |
| GO:0050767 | regulation of neurogenesis | 847 | 84 | 50,07 | 0,0236 |
| GO:0051146 | striated muscle cell differentiation | 318 | 20 | 18,8 | 0,02384 |
| GO:0040011 | locomotion | 1606 | 123 | 94,93 | 0,02416 |
| GO:0050679 | positive regulation of epithelial cell proliferation | 177 | 15 | 10,46 | 0,02435 |
| GO:0009953 | dorsal/ventral pattern formation | 142 | 17 | 8,39 | 0,02469 |
| GO:0051674 | localization of cell | 864 | 65 | 51,07 | 0,02526 |
| GO:0007422 | peripheral nervous system development | 96 | 10 | 5,67 | 0,02561 |
| GO:0060713 | labyrinthine layer morphogenesis | 29 | 5 | 1,71 | 0,02593 |
| GO:0031128 | developmental induction | 43 | 6 | 2,54 | 0,02741 |
| GO:0043550 | regulation of lipid kinase activity | 62 | 7 | 3,66 | 0,02743 |
| GO:2000737 | negative regulation of stem cell differentiation | 20 | 4 | 1,18 | 0,02749 |
| GO:0007565 | female pregnancy | 219 | 22 | 12,94 | 0,02803 |
| GO:0060412 | ventricular septum morphogenesis | 40 | 6 | 2,36 | 0,02878 |
| GO:0006325 | chromatin organization | 731 | 49 | 43,21 | 0,02909 |
| GO:0141193 | nuclear receptor-mediated signaling pathway | 181 | 22 | 10,7 | 0,02936 |
| GO:0045637 | regulation of myeloid cell differentiation | 212 | 18 | 12,53 | 0,02996 |
| GO:0071805 | potassium ion transmembrane transport | 187 | 15 | 11,05 | 0,03021 |
| GO:0048659 | smooth muscle cell proliferation | 147 | 16 | 8,69 | 0,03052 |
| GO:0042633 | hair cycle | 111 | 13 | 6,56 | 0,03059 |
| GO:0072576 | liver morphogenesis | 21 | 4 | 1,24 | 0,03088 |
| GO:0003272 | endocardial cushion formation | 21 | 3 | 1,24 | 0,03095 |
| GO:0032870 | cellular response to hormone stimulus | 766 | 67 | 45,28 | 0,03196 |
| GO:0007157 | heterophilic cell-cell adhesion | 41 | 6 | 2,42 | 0,03209 |

| | | | | | |
|---|---|---|---|---|---|
| GO:0001702 | gastrulation with mouth forming second | 41 | 6 | 2,42 | 0,03209 |
| GO:0086014 | atrial cardiac muscle cell action potential | 21 | 4 | 1,24 | 0,03241 |
| GO:0001889 | liver development | 222 | 22 | 13,12 | 0,03261 |
| GO:0048863 | stem cell differentiation | 227 | 32 | 13,42 | 0,0327 |
| GO:0050772 | positive regulation of axonogenesis | 102 | 8 | 6,03 | 0,0327 |
| GO:0060487 | lung epithelial cell differentiation | 31 | 5 | 1,83 | 0,03369 |
| GO:0010771 | negative regulation of cell morphogenesi... | 88 | 10 | 5,2 | 0,03442 |
| GO:0043507 | positive regulation of JUN kinase activi... | 76 | 9 | 4,49 | 0,03462 |
| GO:0007417 | central nervous system development | 1117 | 98 | 66,02 | 0,03543 |
| GO:1901570 | fatty acid derivative biosynthetic proce... | 74 | 8 | 4,37 | 0,03561 |
| GO:0045907 | positive regulation of vasoconstriction | 42 | 6 | 2,48 | 0,03564 |
| GO:0001756 | somitogenesis | 101 | 11 | 5,97 | 0,03592 |
| GO:0008284 | positive regulation of cell population p... | 845 | 66 | 49,95 | 0,03621 |
| GO:0008285 | negative regulation of cell population p... | 624 | 53 | 36,88 | 0,03629 |
| GO:0001947 | heart looping | 89 | 10 | 5,26 | 0,03682 |
| GO:0071773 | cellular response to BMP stimulus | 170 | 19 | 10,05 | 0,03684 |
| GO:0045596 | negative regulation of cell differentiat... | 679 | 71 | 40,13 | 0,03698 |
| GO:0014033 | neural crest cell differentiation | 100 | 13 | 5,91 | 0,03708 |
| GO:0048705 | skeletal system morphogenesis | 269 | 29 | 15,9 | 0,03734 |
| GO:0031017 | exocrine pancreas development | 22 | 4 | 1,3 | 0,03782 |
| GO:0072079 | nephron tubule formation | 22 | 4 | 1,3 | 0,03782 |
| GO:0072215 | regulation of metanephros development | 22 | 4 | 1,3 | 0,03782 |
| GO:0060259 | regulation of feeding behavior | 32 | 5 | 1,89 | 0,03805 |
| GO:2000036 | regulation of stem cell population maint... | 32 | 5 | 1,89 | 0,03805 |
| GO:0001662 | behavioral fear response | 54 | 7 | 3,19 | 0,03869 |

| | | | | | |
|---|---|---|---|---|---|
| GO:0045471 | response to ethanol | 208 | 19 | 12,29 | 0,03921 |
| GO:0060612 | adipose tissue development | 43 | 6 | 2,54 | 0,03943 |
| GO:0007223 | Wnt signaling pathway, calcium modulatin... | 43 | 6 | 2,54 | 0,03943 |
| GO:0007218 | neuropeptide signaling pathway | 78 | 9 | 4,61 | 0,03998 |
| GO:0034220 | monoatomic ion transmembrane transport | 1117 | 57 | 66,02 | 0,04054 |
| GO:0031323 | regulation of cellular metabolic process | 5298 | 350 | 313,16 | 0,04139 |
| GO:0070301 | cellular response to hydrogen peroxide | 91 | 10 | 5,38 | 0,04194 |
| GO:0071236 | cellular response to antibiotic | 169 | 16 | 9,99 | 0,04211 |
| GO:0006935 | chemotaxis | 532 | 37 | 31,45 | 0,0422 |
| GO:0001892 | embryonic placenta development | 93 | 15 | 5,5 | 0,04249 |
| GO:0042475 | odontogenesis of dentin-containing tooth | 85 | 11 | 5,02 | 0,04253 |
| GO:0046677 | response to antibiotic | 421 | 39 | 24,88 | 0,04256 |
| GO:0048596 | embryonic camera-type eye morphogenesis | 45 | 7 | 2,66 | 0,04341 |
| GO:0007520 | myoblast fusion | 40 | 6 | 2,36 | 0,04354 |
| GO:0060602 | branch elongation of an epithelium | 23 | 4 | 1,36 | 0,04372 |
| GO:0071498 | cellular response to fluid shear stress | 23 | 4 | 1,36 | 0,04372 |
| GO:0031641 | regulation of myelination | 41 | 5 | 2,42 | 0,04373 |
| GO:0001501 | skeletal system development | 556 | 52 | 32,86 | 0,04395 |
| GO:0060008 | Sertoli cell differentiation | 32 | 7 | 1,89 | 0,04426 |
| GO:0002068 | glandular epithelial cell development | 33 | 5 | 1,95 | 0,04449 |
| GO:0031102 | neuron projection regeneration | 70 | 7 | 4,14 | 0,04451 |
| GO:0009200 | deoxyribonucleoside triphosphate metabol... | 23 | 4 | 1,36 | 0,04453 |
| GO:0009746 | response to hexose | 262 | 18 | 15,49 | 0,04454 |
| GO:0048644 | muscle organ morphogenesis | 78 | 7 | 4,61 | 0,04456 |
| GO:0060571 | morphogenesis of an epithelial fold | 24 | 3 | 1,42 | 0,04465 |
| GO:0050680 | negative regulation of epithelial cell p... | 118 | 11 | 6,97 | 0,04596 |
| GO:0060997 | dendritic spine morphogenesis | 63 | 5 | 3,72 | 0,04606 |

| | | | | | |
|---|---|---|---|---|---|
| GO:0042476 | odontogenesis | 128 | 17 | 7,57 | 0,04632 |
| GO:0008219 | cell death | 1862 | 128 | 110,06 | 0,04651 |
| GO:0014031 | mesenchymal cell development | 93 | 10 | 5,5 | 0,04754 |
| GO:0017085 | response to insecticide | 34 | 5 | 2,01 | 0,04778 |
| GO:0021772 | olfactory bulb development | 34 | 5 | 2,01 | 0,04778 |
| GO:0021854 | hypothalamus development | 34 | 5 | 2,01 | 0,04778 |
| GO:0048703 | embryonic viscerocranium morphogenesis | 34 | 5 | 2,01 | 0,04778 |
| GO:0035116 | embryonic hindlimb morphogenesis | 34 | 5 | 2,01 | 0,04778 |
| GO:0043408 | regulation of MAPK cascade | 684 | 58 | 40,43 | 0,04821 |
| GO:0007189 | adenylate cyclase-activating G protein-c... | 94 | 13 | 5,56 | 0,04878 |
| GO:0071300 | cellular response to retinoic acid | 81 | 9 | 4,79 | 0,04903 |
| GO:0014075 | response to amine | 70 | 9 | 4,14 | 0,04963 |
| GO:0035176 | social behavior | 57 | 7 | 3,37 | 0,04971 |
| GO:0051865 | protein autoubiquitination | 57 | 7 | 3,37 | 0,04971 |

**Supplementary Table 10**

| Gene ID | Description | Gene name |
|---|---|---|
| HarAnt_Hap1.g165 | B-cell CLL lymphoma 6a (zinc finger protein 51) | BCL6 |
| HarAnt_Hap1.g201 | Tubulin is the major constituent of microtubules. It binds two moles of GTP, one at an exchangeable site on the beta chain and one at a non-exchangeable site on the alpha chain | TUBB4A |
| HarAnt_Hap1.g348 | Prostaglandin E synthase | PTGES2 |
| HarAnt_Hap1.g509 | acetoacetyl-CoA synthetase | AACS |
| HarAnt_Hap1.g742 | Bone morphogenetic protein receptor | BMPR1B |
| HarAnt_Hap1.g11031 | 7-dehydrocholesterol reductase | DHCR7 |
| HarAnt_Hap1.g11170 | ERO1-like (S. cerevisiae) | ERO1L |
| HarAnt_Hap1.g11253 | Belongs to the TRAFAC class myosin-kinesin ATPase superfamily. Kinesin family | KIF23 |
| HarAnt_Hap1.g11414 | acyl-CoA synthetase family member 3 | ACSF3 |
| HarAnt_Hap1.g15453 | Phospholipase C eta | PLCH2 |
| HarAnt_Hap1.g15533 | Itchy E3 ubiquitin protein ligase | ITCH |
| HarAnt_Hap1.g15598 | Nephronophthisis 4 | NPHP4 |
| HarAnt_Hap1.g15610 | Oxysterol binding protein-like 2b | OSBPL2 |
| HarAnt_Hap1.g16923 | MICAL-like | MICALL2 |
| HarAnt_Hap1.g16994 | Centlein, centrosomal protein | CNTLN |
| HarAnt_Hap1.g17019 | Type II inositol 3,4-bisphosphate | INPP4B |
| HarAnt_Hap1.g17230 | regulatory subunit | PIK3R6 |
| HarAnt_Hap1.g17414 | 3-hydroxybutyrate dehydrogenase, type 2 | BDH2 |
| HarAnt_Hap1.g18372 | Solute carrier family 9, subfamily A (NHE1, cation proton antiporter 1), member 1 | SLC9A1 |
| HarAnt_Hap1.g18497 | ubiquitin-conjugating enzyme | UBE2E1 |
| HarAnt_Hap1.g19011 | Zinc binding alcohol dehydrogenase domain containing 2 | ZADH2 |
| HarAnt_Hap1.g19098 | Abhydrolase domain containing | ABHD5 |
| HarAnt_Hap1.g19316 | Breast carcinoma amplified sequence 3 | BCAS3 |
| HarAnt_Hap1.g19327 | Aldehyde dehydrogenase 3 family, member A2a | ALDH3A2 |
| HarAnt_Hap1.g19581 | Schwannomin interacting protein 1 | SCHIP1 |
| HarAnt_Hap1.g19951 | Dynein assembly factor with WDR repeat domains | DAW1 |
| HarAnt_Hap1.g20305 | Peroxisome biogenesis factor 13 | PEX13 |

| | | |
|---|---|---|
| HarAnt_Hap1.g20461 | Endoplasmic reticulum lectin 1 | ERLEC1 |
| HarAnt_Hap1.g20659 | Pleckstrin | PLEK |
| HarAnt_Hap1.g20790 | Serologically defined colon cancer antigen 8 | SDCCAG8 |
| HarAnt_Hap1.g20796 | F-box protein 5 | FBXO5 |
| HarAnt_Hap1.g20797 | Anaphase promoting complex subunit 1 | ANAPC1 |
| HarAnt_Hap1.g20853 | LIM-domain binding factor 3a | LDB3 |
| HarAnt_Hap1.g21269 | Microtubule associated monooxygenase, calponin and LIM domain containing | MICAL3 |
| HarAnt_Hap1.g21272 | Thromboxane A synthase 1 (platelet, cytochrome P450, family 5, subfamily A) | TBXAS1 |
| HarAnt_Hap1.g21368 | v-Ki-ras2 Kirsten rat sarcoma viral oncogene homolog | KRAS |
| HarAnt_Hap1.g21495 | Nuclear receptor subfamily 2, group F, member 2 | NR2F2 |
| HarAnt_Hap1.g21756 | caldesmon 1 | CALD1 |
| HarAnt_Hap1.g21837 | Cellular retinoic acid binding protein | CRABP1 |
| HarAnt_Hap1.g22833 | Galactosidase, alpha | GLA |
| HarAnt_Hap1.g22945 | Sprouty homolog | SPRY1 |
| HarAnt_Hap1.g22963 | coiled-coil | CCDC69 |
| HarAnt_Hap1.g1941 | pre-B-cell leukemia transcription factor | PBX1 |
| HarAnt_Hap1.g2173 | Kelch-like 20 (Drosophila) | KLHL20 |
| HarAnt_Hap1.g2195 | Fatty acid binding protein 6, ileal (gastrotropin) | FABP6 |
| HarAnt_Hap1.g2369 | Belongs to the profilin family | PFN2 |
| HarAnt_Hap1.g2503 | homolog subfamily B member | DNAJB6 |
| HarAnt_Hap1.g2516 | neuronal | ANK2 |
| HarAnt_Hap1.g2640 | Insulin-induced gene | INSIG2 |
| HarAnt_Hap1.g2811 | Belongs to the TRAFAC class myosin-kinesin ATPase superfamily. Myosin family | MYO1B |
| HarAnt_Hap1.g2877 | erythroblastic leukemia viral oncogene homolog | ERBB4 |
| HarAnt_Hap1.g2997 | Insulin receptor substrate | IRS2 |
| HarAnt_Hap1.g3225 | Nuclear receptor subfamily 0 group B member | NR0B1 |
| HarAnt_Hap1.g3229 | Tripartite motif-containing 13 | TRIM13 |
| HarAnt_Hap1.g3557 | casein kinase | CSNK2A1 |
| HarAnt_Hap1.g3562 | Myotubularin related protein 14 | MTMR14 |
| HarAnt_Hap1.g3668 | Zinc finger protein | ZPR1 |
| HarAnt_Hap1.g3674 | Dehydrogenase reductase SDR family member | DHRS3 |
| HarAnt_Hap1.g3741 | Solute carrier family 16 (monocarboxylate transporter), member 1 | SLC16A1 |

| | | |
|---|---|---|
| HarAnt_Hap1.g4675 | Hydroxyacid oxidase (glycolate oxidase) 1 | HAO1 |
| HarAnt_Hap1.g4753 | intermediate filament bundle assembly | KRT14 |
| HarAnt_Hap1.g4773 | Family with sequence similarity 57, member | FAM57B |
| HarAnt_Hap1.g4776 | CDP-diacylglycerol--inositol 3-phosphatidyltransferase | CDIPT |
| HarAnt_Hap1.g4782 | forkhead box | FOXJ1 |
| HarAnt_Hap1.g4808 | Ectonucleotide pyrophosphatase phosphodiesterase | ENPP7 |
| HarAnt_Hap1.g5000 | Speckle-type POZ | SPOP |
| HarAnt_Hap1.g5225 | SRY (sex determining region Y)-box | SOX9 |
| HarAnt_Hap1.g5502 | Sphingosine kinase | SPHK1 |
| HarAnt_Hap1.g6062 | Thioredoxin-related transmembrane protein | TMX1 |
| HarAnt_Hap1.g6063 | FERM domain containing 6 | FRMD6 |
| HarAnt_Hap1.g6064 | Guanine nucleotide-binding proteins (G proteins) are involved as a modulator or transducer in various transmembrane signaling systems. The beta and gamma chains are required for the GTPase activity, for replacement of GDP by GTP, and for G protein- effector interaction | GNG2 |
| HarAnt_Hap1.g6123 | Lipid phosphate phosphatase-related protein type | PLPPR3 |
| HarAnt_Hap1.g6191 | AHA1, activator of heat shock protein ATPase homolog 1, like | AHSA1 |
| HarAnt_Hap1.g6207 | Zinc finger, FYVE | ZFYVE21 |
| HarAnt_Hap1.g6209 | BTB POZ domain-containing protein | BTBD6 |
| HarAnt_Hap1.g6608 | protein domain specific binding | IPCEF1 |
| HarAnt_Hap1.g6664 | activator of morphogenesis | DAAM2 |
| HarAnt_Hap1.g6693 | Rho-associated, coiled-coil containing protein kinase 2a | ROCK2 |
| HarAnt_Hap1.g6711 | Tubulin is the major constituent of microtubules. It binds two moles of GTP, one at an exchangeable site on the beta chain and one at a non-exchangeable site on the alpha chain | TUBB4A |
| HarAnt_Hap1.g6811 | Responsible for the deiodination of T4 (3,5,3',5'- tetraiodothyronine) | DIO2 |
| HarAnt_Hap1.g7216 | acyl-Coenzyme A binding domain containing 3 | ACBD3 |
| HarAnt_Hap1.g7832 | complement | C3 |
| HarAnt_Hap1.g7835 | Serine incorporator | SERINC2 |
| HarAnt_Hap1.g7896 | Abhydrolase domain containing | ABHD5 |
| HarAnt_Hap1.g7970 | HEPACAM family member 2 | HEPACAM2 |
| HarAnt_Hap1.g8477 | Metastasis suppressor | MTSS1 |
| HarAnt_Hap1.g8736 | Anaphase-promoting complex subunit 4 WD40 domain | FZR1 |
| HarAnt_Hap1.g9020 | Mast stem cell growth factor receptor | KIT |
| HarAnt_Hap1.g9052 | Guanine nucleotide binding protein (G protein), beta polypeptide | GNB4 |

| | | |
|---|---|---|
| HarAnt_Hap1.g9226 | synthase | PTGS2 |
| HarAnt_Hap1.g9280 | Cell surface proteoglycan that bears heparan sulfate | GPC1 |
| HarAnt_Hap1.g9282 | large homolog | dlg1 |
| HarAnt_Hap1.g9392 | regulatory subunit | PIK3R3 |
| HarAnt_Hap1.g9792 | P21 protein (Cdc42 Rac)-activated kinase 3 | PAK3 |
| HarAnt_Hap1.g9997 | Dihydropyrimidinase-like 3 | DPYSL3 |
| HarAnt_Hap1.g10065 | Bromodomain and WD | BRWD3 |
| HarAnt_Hap1.g10172 | Active breakpoint cluster region-related | ABR |
| HarAnt_Hap1.g10195 | Peroxiredoxin 5 | PRDX5 |
| HarAnt_Hap1.g10596 | Sulfotransferase family 4A, member 1 | SULT4A1 |
| HarAnt_Hap1.g10628 | bicaudal D homolog | BICD1 |
| HarAnt_Hap1.g10639 | phytanoyl-CoA | PHYH |
| HarAnt_Hap1.g10653 | Phosphoinositide-3-kinase, catalytic, gamma polypeptide | PIK3CG |
| HarAnt_Hap1.g10723 | tyrosine | TH |
| HarAnt_Hap1.g10850 | ubiquitin-conjugating enzyme | UBE2H |
| HarAnt_Hap1.g11986 | Dolichyl pyrophosphate phosphatase 1 | DOLPP1 |
| HarAnt_Hap1.g12036 | Tubulin tyrosine ligase-like family, member 11 | TTLL11 |
| HarAnt_Hap1.g12057 | Phosphoinositide-3-kinase, regulatory subunit 1 (p85 alpha) | PIK3R1 |
| HarAnt_Hap1.g12451 | Lipid phosphate phosphatase-related protein type | PLPPR1 |
| HarAnt_Hap1.g12474 | Phosphatidylinositol glycan anchor biosynthesis, class O | PIGO |
| HarAnt_Hap1.g12483 | Retinoid X receptor, alpha a | RXRA |
| HarAnt_Hap1.g13084 | Zinc finger, RAN-binding domain containing | ZRANB1 |
| HarAnt_Hap1.g13109 | F-box and leucine-rich repeat protein 15 | FBXL15 |
| HarAnt_Hap1.g13112 | Belongs to the fatty acid desaturase type 1 family | SCD |
| HarAnt_Hap1.g14033 | Signal sequence receptor, alpha | SSR1 |
| HarAnt_Hap1.g14034 | Nebulette | NEBL |
| HarAnt_Hap1.g14041 | TRAF2 and NCK interacting kinase | TNIK |
| HarAnt_Hap1.g14147 | Spindle and centriole associated protein 1 | SPICE1 |
| HarAnt_Hap1.g14395 | ADP-ribosylation factor-like | ARL2 |
| HarAnt_Hap1.g14446 | Belongs to the glycosyl hydrolase 1 family | GBA3 |
| HarAnt_Hap1.g14454 | Tyrosinase-related protein | TYRP1 |

**Supplementary Table 11**

| Gene | OG | pvalue | fdr |
|---|---|---|---|
| HarAnt_Hap1.g10130 | OG0009506 | 0.0107 | 0.234886438356164 |
| HarAnt_Hap1.g10161 | OG0009494 | 0.00443 | 0.145734863013699 |
| HarAnt_Hap1.g10253 | OG0001634 | 0.00119 | 0.0642198876404494 |
| HarAnt_Hap1.g10484 | OG0011405 | 0.01526 | 0.278683574144487 |
| HarAnt_Hap1.g10561 | OG0011384 | 0.00027 | 0.0275917021276596 |
| HarAnt_Hap1.g10589 | OG0011377 | 0.01401 | 0.27300252 |
| HarAnt_Hap1.g10642 | OG0011356 | 0.00046 | 0.036823 |
| HarAnt_Hap1.g10676 | OG0011344 | 0.03273 | 0.388153555555556 |
| HarAnt_Hap1.g10772 | OG0011312 | 0.00265 | 0.102644758064516 |
| HarAnt_Hap1.g1087 | OG0001673 | 0.00229 | 0.0981125641025641 |
| HarAnt_Hap1.g1105 | OG0011240 | 0.02097 | 0.329146764705882 |
| HarAnt_Hap1.g11110 | OG0000739 | 0.00545 | 0.169336538461538 |
| HarAnt_Hap1.g11265 | OG0005564 | 0.02292 | 0.338567300613497 |
| HarAnt_Hap1.g11298 | OG0008769 | 0.00484 | 0.153950463576159 |
| HarAnt_Hap1.g11318 | OG0008777 | 0.02524 | 0.345651794871795 |
| HarAnt_Hap1.g11319 | OG0008778 | 0.03542 | 0.400287670588235 |
| HarAnt_Hap1.g11348 | OG0005572 | 0.01792 | 0.302572264808362 |
| HarAnt_Hap1.g11356 | OG0008796 | 0.02651 | 0.354570498614958 |
| HarAnt_Hap1.g11486 | OG0001575 | 6e-05 | 0.009606 |
| HarAnt_Hap1.g11501 | OG0008845 | 0.02178 | 0.336364437299035 |
| HarAnt_Hap1.g11654 | OG0000841 | 2e-05 | 0.00505578947368421 |
| HarAnt_Hap1.g11700 | OG0000974.ab | 0.02453 | 0.345651794871795 |
| HarAnt_Hap1.g12026 | OG0009182 | 0.01002 | 0.228085592417062 |
| HarAnt_Hap1.g12093 | OG0009202 | 0.01428 | 0.273254342629482 |
| HarAnt_Hap1.g12096 | OG0009203 | 0.00793 | 0.211598833333333 |
| HarAnt_Hap1.g12118 | OG0009215 | 0.0423 | 0.440709110629067 |
| HarAnt_Hap1.g12122 | OG0009219 | 0.03604 | 0.403496783216783 |
| HarAnt_Hap1.g12164 | OG0009226 | 0.0199 | 0.320737248322148 |
| HarAnt_Hap1.g12166 | OG0009228 | 0.0466 | 0.456723407707911 |
| HarAnt_Hap1.g12234 | OG0009245 | 0.01652 | 0.289227927272727 |

| | | | |
|---|---|---|---|
| HarAnt_Hap1.g12333 | OG0009282 | 0.04533 | 0.449169444444444 |
| HarAnt_Hap1.g12339 | OG0009286 | 0.01153 | 0.245250530973451 |
| HarAnt_Hap1.g12352 | OG0005736 | 0.00029 | 0.029018125 |
| HarAnt_Hap1.g12360 | OG0009294 | 0.00255 | 0.0995743902439024 |
| HarAnt_Hap1.g12399 | OG0009308 | 0.03381 | 0.397040171149144 |
| HarAnt_Hap1.g12527 | OG0009345 | 0.03783 | 0.415053767123288 |
| HarAnt_Hap1.g12630 | OG0005682 | 0.00229 | 0.0981125641025641 |
| HarAnt_Hap1.g12855 | OG0006645 | 0.00576 | 0.172908 |
| HarAnt_Hap1.g12870 | OG0009887 | 0.00075 | 0.05003125 |
| HarAnt_Hap1.g12901 | OG0009894 | 0.009 | 0.220512118226601 |
| HarAnt_Hap1.g12957 | OG0001192.aa | 0.03723 | 0.412017718894009 |
| HarAnt_Hap1.g13023 | OG0001016 | 0.01808 | 0.302572264808362 |
| HarAnt_Hap1.g13104 | OG0009947 | 0.00406 | 0.13829914893617 |
| HarAnt_Hap1.g13125 | OG0009955 | 0.04227 | 0.440709110629067 |
| HarAnt_Hap1.g13167 | OG0000957.ab | 0.04316 | 0.442773375796178 |
| HarAnt_Hap1.g13203 | OG0009977 | 0.00255 | 0.0995743902439024 |
| HarAnt_Hap1.g13247 | OG0009988 | 0.04674 | 0.456723407707911 |
| HarAnt_Hap1.g13277 | OG0009997 | 2e-05 | 0.00505578947368421 |
| HarAnt_Hap1.g13297 | OG0010000 | 1e-05 | 0.00369461538461538 |
| HarAnt_Hap1.g13392 | OG0010023 | 0.00236 | 0.0981125641025641 |
| HarAnt_Hap1.g13489 | OG0004013 | 0.01141 | 0.2446528125 |
| HarAnt_Hap1.g13680 | OG0003655 | 0.00041 | 0.0364672222222222 |
| HarAnt_Hap1.g13707 | OG0003870 | 0.00864 | 0.217266596858639 |
| HarAnt_Hap1.g13777 | OG0010868 | 0.0055 | 0.169336538461538 |
| HarAnt_Hap1.g13858 | OG0010891 | 0.04875 | 0.471215855130785 |
| HarAnt_Hap1.g13928 | OG0010905 | 0.00048 | 0.0371845161290323 |
| HarAnt_Hap1.g13973 | OG0010915 | 0.0409 | 0.431742197802198 |
| HarAnt_Hap1.g14024 | OG0010934 | 0.00055 | 0.040025 |
| HarAnt_Hap1.g14097 | OG0004560 | 0.00994 | 0.228085592417062 |
| HarAnt_Hap1.g14144 | OG0010946 | 0.03014 | 0.374089329896907 |
| HarAnt_Hap1.g14514 | OG0006595 | 0.00192 | 0.0899979611650486 |
| HarAnt_Hap1.g15118 | OG0009418 | 0.03022 | 0.374089329896907 |
| HarAnt_Hap1.g15177 | OG0009408 | 0.01001 | 0.228085592417062 |

| | | | |
|---|---|---|---|
| HarAnt_Hap1.g15256 | OG0009388 | 0.00504 | 0.15884431372549 |
| HarAnt_Hap1.g15337 | OG0009373 | 0.03109 | 0.376134181360201 |
| HarAnt_Hap1.g15479 | OG0007257 | 0.02647 | 0.354570498614958 |
| HarAnt_Hap1.g15539 | OG0007269 | 0.03666 | 0.409483674418605 |
| HarAnt_Hap1.g15553 | OG0007274 | 0.03201 | 0.381498833746898 |
| HarAnt_Hap1.g15806 | OG0007362 | 0.03899 | 0.421285560538117 |
| HarAnt_Hap1.g15832 | OG0007368 | 0.02378 | 0.342681107784431 |
| HarAnt_Hap1.g15868 | OG0007378 | 0.03963 | 0.424872522321429 |
| HarAnt_Hap1.g15885 | OG0007384 | 5e-04 | 0.0375234375 |
| HarAnt_Hap1.g15895 | OG0001258 | 0.01154 | 0.245250530973451 |
| HarAnt_Hap1.g15939 | OG0007400 | 0.01976 | 0.320632702702703 |
| HarAnt_Hap1.g16000 | OG0002070 | 0.03765 | 0.414754472477064 |
| HarAnt_Hap1.g16005 | OG0001513 | 0.00338 | 0.118497372262774 |
| HarAnt_Hap1.g16018 | OG0001514 | 0.015 | 0.278683574144487 |
| HarAnt_Hap1.g16105 | OG0007453 | 0 | 0 |
| HarAnt_Hap1.g16218 | OG0007482 | 0.00876 | 0.218001450777202 |
| HarAnt_Hap1.g16227 | OG0007487 | 0.00108 | 0.0596234482758621 |
| HarAnt_Hap1.g16249 | OG0001517 | 0.00921 | 0.220512118226601 |
| HarAnt_Hap1.g16941 | OG0011623 | 0.0149 | 0.278683574144487 |
| HarAnt_Hap1.g17033 | OG0011595 | 0.02168 | 0.335900129032258 |
| HarAnt_Hap1.g17113 | OG0011568 | 0.01525 | 0.278683574144487 |
| HarAnt_Hap1.g17157 | OG0000437.aa | 0.03909 | 0.421285560538117 |
| HarAnt_Hap1.g17222 | OG0000907 | 0.00506 | 0.15884431372549 |
| HarAnt_Hap1.g17479 | OG0011475 | 0.04962 | 0.471757114624506 |
| HarAnt_Hap1.g17616 | OG0011439 | 0.00835 | 0.214618263157895 |
| HarAnt_Hap1.g17670 | OG0011423 | 0.01749 | 0.300015964285714 |
| HarAnt_Hap1.g17742 | OG0000919 | 0.03072 | 0.376134181360201 |
| HarAnt_Hap1.g17757 | OG0006380 | 0.00882 | 0.218363195876289 |
| HarAnt_Hap1.g1798 | OG0000390 | 0.01557 | 0.279039962686567 |
| HarAnt_Hap1.g18355 | OG0007736 | 0.00023 | 0.0251065909090909 |
| HarAnt_Hap1.g18358 | OG0007735 | 0.00055 | 0.040025 |
| HarAnt_Hap1.g18490 | OG0007707 | 0.02796 | 0.362951027027027 |
| HarAnt_Hap1.g18560 | OG0001528 | 6e-05 | 0.009606 |
| HarAnt_Hap1.g18610 | OG0005074 | 0.00295 | 0.110694140625 |

| | | | |
|---|---|---|---|
| HarAnt_Hap1.g1886 | OG0008410 | 0.01232 | 0.259530526315789 |
| HarAnt_Hap1.g19155 | OG0006631 | 0.01515 | 0.278683574144487 |
| HarAnt_Hap1.g19400 | OG0009705 | 0.00842 | 0.214618263157895 |
| HarAnt_Hap1.g19415 | OG0009713 | 0.00125 | 0.0667083333333333 |
| HarAnt_Hap1.g19421 | OG0009716 | 0.02882 | 0.367168328912467 |
| HarAnt_Hap1.g19470 | OG0009729 | 0.00316 | 0.114116390977444 |
| HarAnt_Hap1.g19591 | OG0009764 | 0.00103 | 0.0587661176470588 |
| HarAnt_Hap1.g19600 | OG0009766 | 0.00614 | 0.18203962962963 |
| HarAnt_Hap1.g19618 | OG0009776 | 0.01055 | 0.234590972222222 |
| HarAnt_Hap1.g19663 | OG0009794 | 0.03153 | 0.379808304239402 |
| HarAnt_Hap1.g1967 | OG0008390 | 0.04917 | 0.471757114624506 |
| HarAnt_Hap1.g1969 | OG0001972 | 0.00097 | 0.0568159756097561 |
| HarAnt_Hap1.g19702 | OG0009810 | 0.04708 | 0.457743400809717 |
| HarAnt_Hap1.g19794 | OG0009844 | 0.02261 | 0.338567300613497 |
| HarAnt_Hap1.g19838 | OG0009852 | 0.03579 | 0.403496783216783 |
| HarAnt_Hap1.g19908 | OG0009870 | 0.0105 | 0.23456511627907 |
| HarAnt_Hap1.g19938 | OG0009876 | 0.01136 | 0.2446528125 |
| HarAnt_Hap1.g1994 | OG0008382 | 0.0027 | 0.1037448 |
| HarAnt_Hap1.g20245 | OG0010486 | 0.00335 | 0.118309191176471 |
| HarAnt_Hap1.g20258 | OG0010491 | 0.04688 | 0.456723407707911 |
| HarAnt_Hap1.g20324 | OG0010509 | 0.04106 | 0.432480657894737 |
| HarAnt_Hap1.g20348 | OG0010518 | 0.01065 | 0.234886438356164 |
| HarAnt_Hap1.g20517 | OG0010564 | 0.01583 | 0.281598111111111 |
| HarAnt_Hap1.g2058 | OG0008362 | 0.00664 | 0.189832857142857 |
| HarAnt_Hap1.g20587 | OG0003322 | 0.03197 | 0.381498833746898 |
| HarAnt_Hap1.g20608 | OG0006134 | 0.00046 | 0.036823 |
| HarAnt_Hap1.g20620 | OG0010596 | 0.00044 | 0.036823 |
| HarAnt_Hap1.g20633 | OG0010603 | 0.02351 | 0.342681107784431 |
| HarAnt_Hap1.g2064 | OG0008361 | 0.03977 | 0.425423853006682 |
| HarAnt_Hap1.g20678 | OG0010615 | 0.00924 | 0.220512118226601 |
| HarAnt_Hap1.g2112 | OG0008348 | 0.00574 | 0.172908 |
| HarAnt_Hap1.g21309 | OG0005288 | 0.04966 | 0.471757114624506 |
| HarAnt_Hap1.g21320 | OG0005293 | 0.01071 | 0.234886438356164 |
| HarAnt_Hap1.g21379 | OG0000155.ab | 0.0083 | 0.214618263157895 |

| | | | |
|---|---|---|---|
| HarAnt_Hap1.g21388 | OG0005313 | 0.00679 | 0.190715614035088 |
| HarAnt_Hap1.g21463 | OG0000973.aa | 0.02695 | 0.355606730769231 |
| HarAnt_Hap1.g21482 | OG0005338 | 0.03104 | 0.376134181360201 |
| HarAnt_Hap1.g21564 | OG0005357 | 0.02277 | 0.338567300613497 |
| HarAnt_Hap1.g21638 | OG0008180 | 0.01384 | 0.27300252 |
| HarAnt_Hap1.g21659 | OG0005365 | 0.04687 | 0.456723407707911 |
| HarAnt_Hap1.g21847 | OG0010461 | 2e-05 | 0.00505578947368421 |
| HarAnt_Hap1.g21863 | OG0000980.aa | 0.02358 | 0.342681107784431 |
| HarAnt_Hap1.g21904 | OG0000825.aa | 0.01674 | 0.291312391304348 |
| HarAnt_Hap1.g21926 | OG0003216 | 0.00013 | 0.0173441666666667 |
| HarAnt_Hap1.g22086 | OG0004215 | 0.00239 | 0.0981125641025641 |
| HarAnt_Hap1.g22218 | OG0006919 | 0.0369 | 0.41025625 |
| HarAnt_Hap1.g22280 | OG0010043 | 0.03785 | 0.415053767123288 |
| HarAnt_Hap1.g22358 | OG0002120 | 0.02767 | 0.360159918699187 |
| HarAnt_Hap1.g22416 | OG0010091 | 0.02269 | 0.338567300613497 |
| HarAnt_Hap1.g22424 | OG0010094 | 0.02256 | 0.338567300613497 |
| HarAnt_Hap1.g22493 | OG0000157 | 0 | 0 |
| HarAnt_Hap1.g22552 | OG0010142 | 0.00698 | 0.193785780346821 |
| HarAnt_Hap1.g22785 | OG0010201 | 0.00977 | 0.227792766990291 |
| HarAnt_Hap1.g22822 | OG0010215 | 0.00316 | 0.114116390977444 |
| HarAnt_Hap1.g22905 | OG0010240 | 0.03046 | 0.375988692307692 |
| HarAnt_Hap1.g2504 | OG0005406 | 0.03011 | 0.374089329896907 |
| HarAnt_Hap1.g2836 | OG0004222 | 0.03535 | 0.400287670588235 |
| HarAnt_Hap1.g290 | OG0010984 | 0.0045 | 0.146037162162162 |
| HarAnt_Hap1.g2924 | OG0008586 | 9e-05 | 0.0127138235294118 |
| HarAnt_Hap1.g2956 | OG0008595 | 0.00079 | 0.0505578947368421 |
| HarAnt_Hap1.g2965 | OG0008597 | 0.0083 | 0.214618263157895 |
| HarAnt_Hap1.g3003 | OG0008608 | 0.01303 | 0.269754698275862 |
| HarAnt_Hap1.g3175 | OG0008661 | 0.00021 | 0.024015 |
| HarAnt_Hap1.g3193 | OG0008667 | 0.04545 | 0.449169444444444 |
| HarAnt_Hap1.g3238 | OG0008681 | 0.02874 | 0.367168328912467 |
| HarAnt_Hap1.g3271 | OG0008691 | 0.00899 | 0.220512118226601 |

| | | | |
|---|---|---|---|
| HarAnt_Hap1.g3311 | OG0008695 | 0.03737 | 0.412616344827586 |
| HarAnt_Hap1.g3401 | OG0004113 | 0.02111 | 0.329504512987013 |
| HarAnt_Hap1.g3605 | OG0004870 | 3e-05 | 0.0057636 |
| HarAnt_Hap1.g3743 | OG0006967 | 0.04598 | 0.452208834355828 |
| HarAnt_Hap1.g3777 | OG0006981 | 3e-05 | 0.0057636 |
| HarAnt_Hap1.g390 | OG0011010 | 0.00251 | 0.0995743902439024 |
| HarAnt_Hap1.g3921 | OG0003447 | 0.00422 | 0.142737042253521 |
| HarAnt_Hap1.g4198 | OG0007130 | 0.03483 | 0.400287670588235 |
| HarAnt_Hap1.g4209 | OG0000429.ab | 0.00383 | 0.131396357142857 |
| HarAnt_Hap1.g4214 | OG0004908 | 0.01421 | 0.27300252 |
| HarAnt_Hap1.g4276 | OG0007150 | 0.00983 | 0.228084492753623 |
| HarAnt_Hap1.g4310 | OG0007159 | 0.02748 | 0.359636076294278 |
| HarAnt_Hap1.g4447 | OG0007211 | 0.02906 | 0.368272242744063 |
| HarAnt_Hap1.g4509 | OG0001278 | 0.01782 | 0.302436254416961 |
| HarAnt_Hap1.g456 | OG0001405 | 0.00206 | 0.0924689719626168 |
| HarAnt_Hap1.g4697 | OG0007793 | 0.04007 | 0.427680466666667 |
| HarAnt_Hap1.g4721 | OG0007799 | 8e-04 | 0.0505578947368421 |
| HarAnt_Hap1.g486 | OG0011039 | 0.02189 | 0.3366703514377 |
| HarAnt_Hap1.g4956 | OG0007844 | 0.02526 | 0.345651794871795 |
| HarAnt_Hap1.g4958 | OG0007846 | 0.01911 | 0.314333321917808 |
| HarAnt_Hap1.g4972 | OG0001098.aa | 0.01046 | 0.23456511627907 |
| HarAnt_Hap1.g4994 | OG0007860 | 9e-05 | 0.0127138235294118 |
| HarAnt_Hap1.g5034 | OG0007874 | 0.03165 | 0.379808304239402 |
| HarAnt_Hap1.g5096 | OG0007883 | 0.01402 | 0.27300252 |
| HarAnt_Hap1.g513 | OG0011051 | 0.0226 | 0.338567300613497 |
| HarAnt_Hap1.g5191 | OG0007907 | 0.01095 | 0.239058409090909 |
| HarAnt_Hap1.g525 | OG0011053 | 0.0427 | 0.442414267241379 |
| HarAnt_Hap1.g5321 | OG0007939 | 0.0497 | 0.471757114624506 |
| HarAnt_Hap1.g534 | OG0004579 | 0.0346 | 0.400287670588235 |
| HarAnt_Hap1.g5711 | OG0003652 | 0.03171 | 0.379808304239402 |
| HarAnt_Hap1.g576 | OG0011064 | 0.02718 | 0.35765901369863 |
| HarAnt_Hap1.g5811 | OG0003172 | 0.02605 | 0.35285138028169 |
| HarAnt_Hap1.g586 | OG0011068 | 0.02009 | 0.3222813 |

| | | | |
|---|---|---|---|
| HarAnt_Hap1.g5978 | OG0003073 | 0.02578 | 0.351765170454545 |
| HarAnt_Hap1.g6020 | OG0003506 | 0.02502 | 0.345651794871795 |
| HarAnt_Hap1.g604 | OG0011076 | 0.04326 | 0.442773375796178 |
| HarAnt_Hap1.g6154 | OG0002341 | 0.03848 | 0.417199638826185 |
| HarAnt_Hap1.g6225 | OG0006855 | 0.04319 | 0.442773375796178 |
| HarAnt_Hap1.g6257 | OG0000786.a a | 0.01544 | 0.279005730337079 |
| HarAnt_Hap1.g6366 | OG0006806 | 0.03299 | 0.388948823529412 |
| HarAnt_Hap1.g6471 | OG0006765 | 0.02951 | 0.370320078328982 |
| HarAnt_Hap1.g65 | OG0008961 | 0.00664 | 0.189832857142857 |
| HarAnt_Hap1.g6518 | OG0006747 | 0.02659 | 0.354570498614958 |
| HarAnt_Hap1.g6525 | OG0001486 | 0.02593 | 0.352809603399433 |
| HarAnt_Hap1.g6609 | OG0006725 | 0.02524 | 0.345651794871795 |
| HarAnt_Hap1.g6659 | OG0006709 | 0.03827 | 0.417199638826185 |
| HarAnt_Hap1.g6774 | OG0004779 | 0.00308 | 0.112925496183206 |
| HarAnt_Hap1.g6970 | OG0010690 | 0.01764 | 0.300442978723404 |
| HarAnt_Hap1.g7015 | OG0010704 | 0.00218 | 0.0969494444444444 |
| HarAnt_Hap1.g7073 | OG0010724 | 0.0141 | 0.27300252 |
| HarAnt_Hap1.g7075 | OG0010725 | 0.00816 | 0.214618263157895 |
| HarAnt_Hap1.g7114 | OG0010740 | 0.00193 | 0.0899979611650486 |
| HarAnt_Hap1.g7158 | OG0000457.a b | 0.04028 | 0.427816887417219 |
| HarAnt_Hap1.g7257 | OG0010789 | 0.02291 | 0.338567300613497 |
| HarAnt_Hap1.g7284 | OG0010795 | 0.03518 | 0.400287670588235 |
| HarAnt_Hap1.g7297 | OG0004526 | 0.00238 | 0.0981125641025641 |
| HarAnt_Hap1.g7421 | OG0010832 | 0.00248 | 0.0995743902439024 |
| HarAnt_Hap1.g7825 | OG0000885.a b | 0.01653 | 0.289227927272727 |
| HarAnt_Hap1.g7866 | OG0010449 | 0.00073 | 0.0493829577464789 |
| HarAnt_Hap1.g7970 | OG0010422 | 0.00161 | 0.0805503125 |
| HarAnt_Hap1.g8061 | OG0010388 | 0.0019 | 0.0899979611650486 |
| HarAnt_Hap1.g8076 | OG0010379 | 0.03542 | 0.400287670588235 |
| HarAnt_Hap1.g8151 | OG0001306 | 0.01863 | 0.309379448275862 |
| HarAnt_Hap1.g8177 | OG0006044 | 0.02274 | 0.338567300613497 |
| HarAnt_Hap1.g8237 | OG0010326 | 0.00457 | 0.1463314 |

| | | | |
|---|---|---|---|
| HarAnt_Hap1.g830 | OG0011145 | 0.02659 | 0.354570498614958 |
| HarAnt_Hap1.g8373 | OG0002562 | 0.04035 | 0.427816887417219 |
| HarAnt_Hap1.g8514 | OG0006022 | 0.0111 | 0.24015 |
| HarAnt_Hap1.g859 | OG0011159 | 5e-04 | 0.0375234375 |
| HarAnt_Hap1.g8846 | OG0009131 | 0.02013 | 0.3222813 |
| HarAnt_Hap1.g893 | OG0011174 | 0.01372 | 0.27300252 |
| HarAnt_Hap1.g8947 | OG0009096 | 0.03912 | 0.421285560538117 |
| HarAnt_Hap1.g8971 | OG0009089 | 4e-04 | 0.0362490566037736 |
| HarAnt_Hap1.g8977 | OG0009088 | 8e-05 | 0.0123948387096774 |
| HarAnt_Hap1.g899 | OG0011175 | 0.02415 | 0.343741331360947 |
| HarAnt_Hap1.g9020 | OG0009074 | 0.01302 | 0.269754698275862 |
| HarAnt_Hap1.g9040 | OG0009065 | 0.01551 | 0.279005730337079 |
| HarAnt_Hap1.g9261 | OG0009008 | 0.03166 | 0.379808304239402 |
| HarAnt_Hap1.g9426 | OG0004281 | 0.00738 | 0.201398522727273 |
| HarAnt_Hap1.g943 | OG0006318 | 0.00588 | 0.175413913043478 |
| HarAnt_Hap1.g946 | OG0011186 | 0.00849 | 0.214618263157895 |
| HarAnt_Hap1.g962 | OG0005765 | 0.00736 | 0.201398522727273 |
| HarAnt_Hap1.g9633 | OG0009651 | 0.00108 | 0.0596234482758621 |
| HarAnt_Hap1.g964 | OG0011192 | 0.02924 | 0.368608188976378 |
| HarAnt_Hap1.g9672 | OG0009639 | 0.01549 | 0.279005730337079 |
| HarAnt_Hap1.g9706 | OG0009629 | 0.00046 | 0.036823 |
| HarAnt_Hap1.g9804 | OG0000437.ab | 3e-05 | 0.0057636 |
| HarAnt_Hap1.g9811 | OG0009588 | 0.01392 | 0.27300252 |
| HarAnt_Hap1.g9929 | OG0009564 | 0.04961 | 0.471757114624506 |
| HarAnt_Hap1.g9952 | OG0005838 | 0.01619 | 0.286939372693727 |
| HarAnt_Hap1.g996 | OG0011206 | 0.02476 | 0.345651794871795 |
| HarAnt_Hap1.g997 | OG0011207 | 2e-04 | 0.0234292682926829 |
| HarAnt_Hap1.g9985 | OG0009549 | 0.00785 | 0.210634357541899 |

### Supplementary Table 12

| GO.ID | Term | Annotated | Significant | Expected | classicFisher |
|---|---|---|---|---|---|
| **GO:0043524** | negative regulation of neuron apoptotic process | 55 | 12 | 3,83 | 0,00029 |
| **GO:0010501** | RNA secondary structure unwinding | 13 | 5 | 0,9 | 0,00127 |
| **GO:0030595** | leukocyte chemotaxis | 28 | 4 | 1,95 | 0,00302 |
| **GO:0003407** | neural retina development | 33 | 7 | 2,3 | 0,00307 |
| **GO:0071353** | cellular response to interleukin-4 | 10 | 4 | 0,7 | 0,00343 |
| **GO:0014031** | mesenchymal cell development | 23 | 6 | 1,6 | 0,00393 |
| **GO:0071902** | positive regulation of protein serine/threonine kinase activity | 81 | 11 | 5,64 | 0,00614 |
| **GO:0035886** | vascular associated smooth muscle cell differentiation | 12 | 4 | 0,83 | 0,00724 |
| **GO:0014032** | neural crest cell development | 23 | 5 | 1,6 | 0,00934 |
| **GO:0055123** | digestive system development | 58 | 4 | 4,04 | 0,00972 |
| **GO:0032092** | positive regulation of protein binding | 28 | 6 | 1,95 | 0,01094 |
| **GO:0006690** | icosanoid metabolic process | 22 | 4 | 1,53 | 0,01372 |
| **GO:0046031** | ADP metabolic process | 28 | 3 | 1,95 | 0,01388 |
| **GO:1903313** | positive regulation of mRNA metabolic process | 30 | 3 | 2,09 | 0,0139 |
| **GO:0031503** | protein-containing complex localization | 47 | 8 | 3,27 | 0,01434 |
| **GO:0019221** | cytokine-mediated signaling pathway | 126 | 14 | 8,77 | 0,01545 |
| **GO:0007030** | Golgi organization | 39 | 7 | 2,71 | 0,01629 |
| **GO:0032496** | response to lipopolysaccharide | 80 | 11 | 5,57 | 0,02058 |
| **GO:0007596** | blood coagulation | 73 | 9 | 5,08 | 0,02078 |
| **GO:0045665** | negative regulation of neuron differentiation | 61 | 9 | 4,24 | 0,02352 |
| **GO:0046425** | regulation of receptor signaling pathway via JAK-STAT | 30 | 6 | 2,09 | 0,02588 |
| **GO:0006220** | pyrimidine nucleotide metabolic process | 16 | 4 | 1,11 | 0,02609 |
| **GO:0042310** | vasoconstriction | 21 | 4 | 1,46 | 0,02616 |
| **GO:0006361** | transcription initiation at RNA polymerase I promoter | 17 | 4 | 1,18 | 0,02646 |
| **GO:0030521** | androgen receptor signaling pathway | 17 | 4 | 1,18 | 0,02646 |
| **GO:0051603** | obsolete proteolysis involved in protein catabolic process | 174 | 16 | 12,11 | 0,02717 |
| **GO:1901984** | negative regulation of protein acetylation | 10 | 3 | 0,7 | 0,02768 |
| **GO:0001910** | regulation of leukocyte mediated cytotoxicity | 10 | 3 | 0,7 | 0,02768 |
| **GO:0009303** | rRNA transcription | 10 | 3 | 0,7 | 0,02768 |
| **GO:1903727** | positive regulation of phospholipid metabolic process | 10 | 3 | 0,7 | 0,02768 |
| **GO:0051260** | protein homooligomerization | 107 | 13 | 7,44 | 0,03246 |
| **GO:0006084** | acetyl-CoA metabolic process | 11 | 3 | 0,77 | 0,03615 |
| **GO:0032481** | positive regulation of type I interferon production | 11 | 3 | 0,77 | 0,03615 |

| | | | | | |
|---|---|---|---|---|---|
| **GO:0047496** | vesicle transport along microtubule | 11 | 3 | 0,77 | 0,03615 |
| **GO:0000027** | ribosomal large subunit assembly | 11 | 3 | 0,77 | 0,03615 |
| **GO:0010039** | response to iron ion | 11 | 3 | 0,77 | 0,03615 |
| **GO:0000460** | maturation of 5.8S rRNA | 11 | 3 | 0,77 | 0,03615 |
| **GO:0045601** | regulation of endothelial cell differentiation | 11 | 3 | 0,77 | 0,03615 |
| **GO:0045089** | positive regulation of innate immune response | 62 | 4 | 4,31 | 0,03715 |
| **GO:0006364** | rRNA processing | 63 | 10 | 4,38 | 0,03854 |
| **GO:0045840** | positive regulation of mitotic nuclear division | 19 | 4 | 1,32 | 0,03864 |
| **GO:0043065** | positive regulation of apoptotic process | 160 | 15 | 11,13 | 0,03867 |
| **GO:0048732** | gland development | 153 | 13 | 10,64 | 0,03879 |
| **GO:0050768** | negative regulation of neurogenesis | 78 | 11 | 5,43 | 0,04011 |
| **GO:0070301** | cellular response to hydrogen peroxide | 28 | 5 | 1,95 | 0,04111 |
| **GO:1903047** | mitotic cell cycle process | 204 | 16 | 14,19 | 0,04184 |
| **GO:1902680** | positive regulation of RNA biosynthetic process | 395 | 36 | 27,48 | 0,04262 |
| **GO:0010605** | negative regulation of macromolecule metabolic process | 620 | 42 | 43,14 | 0,04297 |
| **GO:0006509** | membrane protein ectodomain proteolysis | 12 | 3 | 0,83 | 0,04578 |
| **GO:0051150** | regulation of smooth muscle cell differentiation | 12 | 3 | 0,83 | 0,04578 |
| **GO:0045824** | negative regulation of innate immune response | 12 | 3 | 0,83 | 0,04578 |
| **GO:0046427** | positive regulation of receptor signaling pathway via JAK-STAT | 12 | 3 | 0,83 | 0,04578 |
| **GO:0002474** | antigen processing and presentation of peptide antigen via MHC class I | 12 | 3 | 0,83 | 0,04578 |
| **GO:0008217** | regulation of blood pressure | 45 | 5 | 3,13 | 0,04592 |

**Supplementary Table 13**

| Hap1 | | | |
|---|---|---|---|
| **Chr** | **Gene name** | **Status** | **Gene ID** |
| SUPER_5 | tryp1 | complete | Hap1.G00000000001 |
| SUPER_5 | tryp1 | complete | Hap1.G00000000002 |
| SUPER_5 | tryp1 | complete | Hap1.G00000000003 |
| SUPER_5 | tryp1 | complete | Hap1.G00000000004 |
| SUPER_5 | tryp1 | complete | Hap1.G00000000005 |
| SUPER_5 | tryp1 | complete | Hap1.G00000000006 |
| SUPER_5 | tryp1 | partial | Hap1.G00000000007 |
| SUPER_5 | tryp1 | complete | Hap1.G00000000008 |
| SUPER_5 | tryp1 | partial | Hap1.G00000000009 |
| SUPER_5 | tryp1 | complete | Hap1.G00000000010 |
| SUPER_5 | tryp3 | complete | Hap1.G00000000011 |
| SUPER_5 | tlp | complete | Hap1.G00000000012 |
| SUPER_5 | afgp1 | partial | Hap1.G00000000013 |
| SUPER_5 | afgp2 | complete | Hap1.G00000000014 |
| SUPER_5 | tryp3 | partial | Hap1.G00000000015 |
| SUPER_5 | tryp3 | partial | Hap1.G00000000016 |
| SUPER_5 | afgp3 | complete | Hap1.G00000000017 |
| SUPER_5 | afgp4 | partial | Hap1.G00000000018 |
| SUPER_5 | tryp3 | partial | Hap1.G00000000019 |
| SUPER_5 | tryp3 | partial | Hap1.G00000000020 |
| SUPER_5 | afgp5 | complete | Hap1.G00000000021 |

| | | | |
|---|---|---|---|
| SUPER_5 | afgp6 | complete | Hap1.G00000000022 |
| SUPER_5 | tryp3 | partial | Hap1.G00000000023 |
| SUPER_5 | tryp3 | partial | Hap1.G00000000024 |
| SUPER_5 | afgp7 | complete | Hap1.G00000000025 |
| SUPER_5 | afgp8 | complete | Hap1.G00000000026 |
| SUPER_5 | tryp3 | partial | Hap1.G00000000027 |
| SUPER_5 | tryp3 | partial | Hap1.G00000000028 |
| SUPER_5 | afgp9 | complete | Hap1.G00000000029 |
| SUPER_5 | afgp10 | complete | Hap1.G00000000030 |
| SUPER_5 | tryp3 | partial | Hap1.G00000000031 |
| SUPER_5 | tryp3 | partial | Hap1.G00000000032 |
| SUPER_5 | afgp11 | complete | Hap1.G00000000033 |
| SUPER_5 | afgp12 | complete | Hap1.G00000000034 |
| SUPER_5 | tryp3 | partial | Hap1.G00000000035 |
| SUPER_5 | tryp3 | partial | Hap1.G00000000036 |
| SUPER_5 | afgp13 | complete | Hap1.G00000000037 |
| SUPER_5 | afgp14 | complete | Hap1.G00000000038 |
| SUPER_5 | tryp3 | partial | Hap1.G00000000039 |
| SUPER_5 | afgp15 | complete | Hap1.G00000000040 |

| | | | |
|---|---|---|---|
| SUPER_5 | afgp/tlp1 | complete | Hap1.G00000000041 |
| SUPER_5 | afgp16 | complete | Hap1.G00000000042 |
| SUPER_5 | afgp17 | partial | Hap1.G00000000043 |
| SUPER_5 | tryp3 | complete | Hap1.G00000000044 |
| SUPER_5 | afgp/tlp2 | partial | Hap1.G00000000045 |
| SUPER_5 | tomm40 | complete | Hap1.G00000000046 |

| **Hap2** | | | |
|---|---|---|---|
| **Chr** | **Gene name** | **Status** | **Gene ID** |
| SUPER_5 | tryp1 | complete | Hap2.G00000000001 |
| SUPER_5 | tryp1 | complete | Hap2.G00000000002 |
| SUPER_5 | tryp1 | complete | Hap2.G00000000003 |
| SUPER_5 | tryp1 | complete | Hap2.G00000000004 |
| SUPER_5 | tryp1 | complete | Hap2.G00000000005 |
| SUPER_5 | tryp1 | complete | Hap2.G00000000006 |
| SUPER_5 | tryp1 | partial | Hap2.G00000000007 |
| SUPER_5 | tryp1 | complete | Hap2.G00000000008 |
| SUPER_5 | tryp1 | partial | Hap2.G00000000009 |
| SUPER_5 | tryp1 | complete | Hap2.G00000000010 |
| SUPER_5 | tryp1 | partial | Hap2.G00000000011 |
| SUPER_5 | tryp1 | complete | Hap2.G00000000012 |
| SUPER_5 | tryp3 | complete | Hap2.G00000000013 |
| SUPER_5 | tlp | complete | Hap2.G00000000014 |
| SUPER_5 | afgp1 | partial | Hap2.G00000000015 |

| | | | |
|---|---|---|---|
| SUPER_5 | afgp2 | complete | Hap2.G00000000016 |
| SUPER_5 | tryp3 | partial | Hap2.G00000000017 |
| SUPER_5 | tryp3 | partial | Hap2.G00000000018 |
| SUPER_5 | afgp3 | complete | Hap2.G00000000019 |
| SUPER_5 | afgp4 | complete | Hap2.G00000000020 |
| SUPER_5 | tryp3 | partial | Hap2.G00000000021 |
| SUPER_5 | tryp3 | partial | Hap2.G00000000022 |
| SUPER_5 | afgp5 | complete | Hap2.G00000000023 |
| SUPER_5 | tryp3 | partial | Hap2.G00000000024 |
| SUPER_5 | afgp7 | complete | Hap2.G00000000025 |
| SUPER_5 | afgp8 | complete | Hap2.G00000000026 |
| SUPER_5 | tryp3 | partial | Hap2.G00000000027 |
| SUPER_5 | tryp3 | partial | Hap2.G00000000028 |
| SUPER_5 | afgp9 | complete | Hap2.G00000000029 |
| SUPER_5 | afgp10 | complete | Hap2.G00000000030 |
| SUPER_5 | tryp3 | partial | Hap2.G00000000031 |
| SUPER_5 | tryp3 | partial | Hap2.G00000000032 |
| SUPER_5 | afgp11 | complete | Hap2.G00000000033 |
| SUPER_5 | afgp12 | complete | Hap2.G00000000034 |
| SUPER_5 | tryp3 | partial | Hap2.G00000000035 |
| SUPER_5 | tryp3 | partial | Hap2.G00000000036 |
| SUPER_5 | afgp13 | complete | Hap2.G00000000037 |
| SUPER_5 | afgp14 | complete | Hap2.G00000000038 |
| SUPER_5 | tryp3 | partial | Hap2.G00000000039 |
| SUPER_5 | afgp15 | complete | Hap2.G00000000040 |
| SUPER_5 | afgp/tlp | complete | Hap2.G00000000041 |
| SUPER_5 | afgp16 | complete | Hap2.G00000000042 |
| SUPER_5 | afgp17 | partial | Hap2.G00000000043 |
| SUPER_5 | tryp3 | complete | Hap2.G00000000044 |
| SUPER_5 | afgp/tlp | complete | Hap2.G00000000045 |
| SUPER_5 | tomm40 | complete | Hap2.G00000000046 |
